# Computational modeling reveals frequency modulation of calcium-cAMP/PKA pathway in dendritic spines

**DOI:** 10.1101/521740

**Authors:** D. Ohadi, D. L. Schmitt, B. Calabrese, S. Halpain, J. Zhang, P. Rangamani

## Abstract

Dendritic spines are the primary excitatory postsynaptic sites that act as subcompartments of signaling. Ca^2+^ is often the first and most rapid signal in spines. Downstream of calcium, the cAMP/PKA pathway plays a critical role in the regulation of spine formation, morphological modifications, and ultimately, learning and memory. While the dynamics of calcium are reasonably well-studied, calcium-induced cAMP/PKA dynamics, particularly with respect to frequency modulation, are not fully explored. In this study, we present a well-mixed model for the dynamics of calcium-induced cAMP/PKA dynamics in dendritic spines. The model is constrained using experimental observations in the literature. Further, we measured the calcium oscillation frequency in dendritic spines of cultured hippocampal CA1 neurons and used these dynamics as model inputs. Our model predicts that the various steps in this pathway act as frequency modulators for calcium and the high frequency of calcium input is filtered by AC1 and PDEs in this pathway such that cAMP/PKA only responds to lower frequencies. This prediction has important implications for noise filtering and long-timescale signal transduction in dendritic spines. A companion manuscript presents a three-dimensional spatial model for the same pathway.

**Statement of Significance:** cAMP/PKA activity triggered by calcium is an essential biochemical pathway for synaptic plasticity, regulating spine structure, and long-term potentiation. In the current study, we predicted that for a given calcium input, AC1, and PDE1 kinetics reflect both the high and the low frequencies with different amplitudes and cAMP/PKA acts as a leaky integrator of calcium because of frequency attenuation by the intermediary steps. These findings have implications for cAMP/PKA signaling in dendritic spines in particular and neuronal signal transduction in general.

## Introduction

Influence of Ca^2+^ in response to neurotransmitter-mediated ion channel opening is one of the first steps in synaptic plasticity [1, 2]. Transient high calcium elevation in postsynaptic domains results in the activation of multiple protein kinases including calcium-calmodulin dependent kinase II (CaMKII) and cAMP-dependent protein kinase A (PKA) among others [3–8]. These protein kinases are associated with long-term potentiation (LTP), memory formation, consolidation, and retrieval and are known to have distinct roles in various aspects of these processes (See Table 1 in Giese and Mizuno [9]).

**Table 1:** cAMP-PKA pathway reactions, reaction types, and reaction rates used in the model

|  | List of Reactions | Reaction Type | Reaction Rate |
| --- | --- | --- | --- |
| Module 1: | Ca<sup>2+</sup>/CaM Complex Formation |  |  |
| 1 | 2 Ca<sup>2+</sup> + CaM $\xrightleftharpoons[k_{b1}]{k_{f1}}$ Ca<sub>2</sub>·CaM | MA<sup>(1)</sup> | $R_1 = k_{f1}[Ca^{2+}]^2[CaM] - k_{b1}[Ca_2CaM]$ |
| 2 | 2 Ca<sup>2+</sup> + Ca<sub>2</sub>·CaM $\xrightleftharpoons[k_{b2}]{k_{f2}}$ Ca<sub>4</sub>·CaM | MA | $R_2 = k_{f2}[Ca^{2+}]^2[Ca_2CaM] - k_{b2}[Ca_4CaM]$ |
| Module 2: | AC1 Activation By Ca<sup>2+</sup>/CaM Complex |  |  |
| 3 | AC1 + Ca<sub>2</sub>·CaM $\xrightleftharpoons[k_{b3}]{k_{f3}}$ AC1·Ca<sub>2</sub>·CaM | MA | $R_3 = k_{f3}[AC1][Ca_2CaM] - k_{b3}[AC1Ca_2CaM]$ |
| 4 | 2 Ca<sup>2+</sup> + AC1·Ca<sub>2</sub>·CaM $\xrightleftharpoons[k_{b4}]{k_{f4}}$ AC1·Ca<sub>4</sub>·CaM | CB<sup>(2)</sup> | $R_4 = \frac{k_{f4}[AC1Ca_2CaM][Ca^{2+}]}{K_{m4} + [Ca^{2+}]} - k_{b4}[AC1Ca_4CaM]$ |
| Module 3: | PDE1 Activation By Ca<sup>2+</sup>/CaM Complex |  |  |
| 5 | PDE1 + Ca<sub>2</sub>·CaM $\xrightleftharpoons[k_{b5}]{k_{f5}}$ PDE1·Ca<sub>2</sub>CaM | MA | $R_5 = k_{f5}[PDE1][Ca_2CaM] - k_{b5}[PDE1Ca_2CaM]$ |
| 6 | 2 Ca<sup>2+</sup> + PDE1·Ca<sub>2</sub>·CaM $\xrightleftharpoons[k_{b6}]{k_{f6}}$ PDE1·Ca<sub>4</sub>·CaM | CB | $R_6 = \frac{k_{f6}[PDE1Ca_2CaM][Ca^{2+}]}{K_{m6} + [Ca^{2+}]} - k_{b6}[PDE1Ca_4CaM]$ |
| Module 4: | cAMP Production |  |  |
| 7 | ATP + **AC1·Ca<sub>4</sub>·CaM**<sup>(3)</sup> $\xrightarrow{K_{cat7}}$ cAMP | MM<sup>(4)</sup> | $R_7 = \frac{K_{cat7}[AC1Ca_4M][ATP]}{K_{m7} + [ATP]} - K_{r7}[cAMP]$ |
| Module 5: | PKA Activation |  |  |
| 8 | R<sub>2</sub>C<sub>2</sub> + 2 cAMP $\xrightleftharpoons[k_{b8}]{k_{f8}}$ R<sub>2</sub>C<sub>2</sub>·cAMP<sub>2</sub> | MA | $R_8 = k_{f8}[R_2C_2][cAMP]^2 - k_{b8}[R_2C_2cAMP_2]$ |
| 9 | R<sub>2</sub>C<sub>2</sub>·cAMP<sub>2</sub> + 2 cAMP $\xrightleftharpoons[k_{b9}]{k_{f9}}$ R<sub>2</sub>C<sub>2</sub>·cAMP<sub>4</sub> | MA | $R_9 = k_{f9}[R_2C_2cAMP_2][cAMP]^2 - k_{b9}[R_2C_2cAMP_4]$ |
| 10 | R<sub>2</sub>C<sub>2</sub>·cAMP<sub>4</sub> $\xrightleftharpoons[k_{b10}]{k_{f10}}$ 2 PKAc + R<sub>2</sub>·cAMP<sub>4</sub> | MA | $R_{10} = k_{f10}[R_2C_2cAMP_4] - k_{b10}[PKAc]^2[R_2cAMP_4]$ |
| Module 6: | PDE Phosphorylation By PKA |  |  |
| 11 | PDE1 + **PKAc** $\xrightarrow{K_{cat11}}$ PDE1P | MM | $R_{11} = \frac{K_{cat11}[PDE1][PKAc]}{K_{m11} + [PDE1]} - K_{r11}[PDE1P]$ |
| 12 | PDE4 + **PKAc** $\xrightarrow{K_{cat12}}$ PDE4P | MM | $R_{12} = \frac{K_{cat12}[PDE4][PKAc]}{K_{m12} + [PDE4]} - K_{r12}[PDE4P]$ |
| Module 7: | cAMP Degradation |  |  |
| 13 | cAMP + **PDE1·Ca<sub>4</sub>·CaM** $\xrightarrow{K_{cat13}}$ AMP | MM | $R_{13} = \frac{K_{cat13}[PDE1Ca_4CaM][cAMP]}{K_{m13} + [cAMP]} - K_{r13}[AMP]$ |
| 14 | cAMP + **PDE4** $\xrightarrow{K_{cat14}}$ AMP | MM | $R_{14} = \frac{K_{cat14}[PDE4][cAMP]}{K_{m14} + [cAMP]} - K_{r14}[AMP]$ |
| 15 | cAMP + **PDE4P** $\xrightarrow{K_{cat15}}$ AMP | MM | $R_{15} = \frac{K_{cat15}[PDE4P][cAMP]}{K_{m15} + [cAMP]} - K_{r15}[AMP]$ |
| Module 8: | AC1 Activation and cAMP Production By G<sub>s</sub> |  |  |
| 16 | ISO + β<sub>2</sub>AR $\xrightleftharpoons[k_{b16}]{k_{f16}}$ ISO·β<sub>2</sub>AR | MA | $R_{16} = k_{f16}[ISO][\beta_2AR] - k_{b16}[ISO \cdot \beta_2AR]$ |
| 17 | ISO·β<sub>2</sub>AR + G<sub>sαβγ</sub> $\xrightleftharpoons[k_{b17}]{k_{f17}}$ ISO·β<sub>2</sub>AR·G<sub>sαβγ</sub> | MA | $R_{17} = k_{f17}[ISO \cdot \beta_2AR][G_{s\alpha\beta\gamma}] - k_{b17}[ISO \cdot \beta_2AR \cdot G_{s\alpha\beta\gamma}]$ |
| 18 | ISO·β<sub>2</sub>AR·G<sub>sαβγ</sub> $\xrightarrow{k_{f18}}$ ISO·β<sub>2</sub>AR·G·βγ + G<sub>sα</sub>GTP | MA | $R_{18} = k_{f18}[ISO \cdot \beta_2AR \cdot G_{s\alpha\beta\gamma}]$ |
| 19 | β<sub>2</sub>AR + G<sub>sαβγ</sub> $\xrightleftharpoons[k_{b19}]{k_{f19}}$ β<sub>2</sub>AR·G<sub>sαβγ</sub> | MA | $R_{19} = k_{f19}[\beta_2AR][G_{s\alpha\beta\gamma}] - k_{b19}[\beta_2AR \cdot G_{s\alpha\beta\gamma}]$ |
| 20 | ISO + β<sub>2</sub>AR·G<sub>sαβγ</sub> $\xrightleftharpoons[k_{b20}]{k_{f20}}$ ISO·β<sub>2</sub>AR·G<sub>sαβγ</sub> | MA | $R_{20} = k_{f20}[ISO][\beta_2AR \cdot G_{s\alpha\beta\gamma}] - k_{b20}[ISO \cdot \beta_2AR \cdot G_{s\alpha\beta\gamma}]$ |
| 21 | ISO·β<sub>2</sub>AR·G<sub>sαβγ</sub> $\xrightarrow{k_{f21}}$ ISO·β<sub>2</sub>AR·G·βγ + G<sub>sα</sub>GTP | MA | $R_{21} = k_{f21}[ISO \cdot \beta_2AR \cdot G_{s\alpha\beta\gamma}]$ |
| 22 | ISO·β<sub>2</sub>AR·G·βγ $\xrightarrow{k_{f22}}$ ISO·β<sub>2</sub>AR + G·βγ | MA | $R_{22} = k_{f22}[ISO \cdot \beta_2AR \cdot G_{\beta\gamma}]$ |
| 23 | G<sub>sα</sub>GTP $\xrightarrow{k_{f23}}$ G<sub>sα</sub>GDP | MA | $R_{23} = k_{f23}[G_{s\alpha\beta\gamma}GTP]$ |
| 24 | G<sub>sα</sub>GDP + G·βγ $\xrightarrow{k_{f24}}$ G<sub>sαβγ</sub> | MA | $R_{24} = k_{f24}[G_{s\alpha\beta\gamma}GDP][G_{\beta\gamma}]$ |
| 25 | AC1 + G<sub>sα</sub>GTP $\xrightleftharpoons[k_{b25}]{k_{f25}}$ AC1·G<sub>sα</sub>GTP | MA | $R_{25} = k_{f25}[AC1][G_{s\alpha}GTP] - k_{b25}[AC1 \cdot G_{s\alpha}GTP]$ |
| 26 | AC1·G<sub>sα</sub>GTP + Ca<sub>4</sub>CaM $\xrightleftharpoons[k_{b26}]{k_{f26}}$ AC1·G<sub>sα</sub>GTP·Ca<sub>4</sub>CaM | MA | $R_{26} = k_{f26}[AC1 \cdot G_{s\alpha}GTP][Ca_4CaM] - k_{b26}[AC1 \cdot G_{s\alpha}GTP \cdot Ca_4CaM]$ |
| 27 | ATP + **AC1·G<sub>sα</sub>GTP·Ca<sub>4</sub>CaM**<sup>(2)</sup> $\xrightarrow{K_{cat27}}$ cAMP | MM | $R_{27} = \frac{K_{cat27}[AC1 \cdot G_{s\alpha}GTP \cdot Ca_4CaM][cAMP]}{K_{m27} + [cAMP]}$ |

(<sup>1</sup>) MA: Mass Action; (<sup>2</sup>) CB: Cooperative Binding; (<sup>3</sup>) Enzyme is in bold type; (<sup>4</sup>) MM: Michaelis-Menten

The dynamics of the protein kinases associated with LTP have been studied using systems biology approaches for more than two decades now [4, 10–12]. In 1999, a seminal paper by Bhalla and Iyengar [13] highlighted the role of coupled signaling between PKA, CaMKII, and MAPK and how they govern the emergent properties of signaling in neurons. Subsequently, Hayer and Bhalla [14] expanded this model to include trafficking of AMPA receprors into the synapse and showed that the protein kinase activity and signaling crosstalk can affect trafficking dynamics. These models are examples of comprehensive pathway maps that account for the extensive detailed interactions between various biochemical components in neurons. Separately, detailed models for specific pathway components have also been developed. For example, the dynamics of CaMKII as a function of calcium influx have been well studied both experimentally [15–17] and theoretically [18, 19]. These studies have established the role of CaMKII as a critical molecule in synaptic plasticity and the role played by CaMKII bistability in determining LTP versus LTD [20–22].

Tightly coupled with calcium dynamics, through various signaling pathways, are the dynamics of another important second messenger, cAMP [23, 24]. Two types of enzymes play important roles in driving parallel Ca^2+^ and cAMP oscillations: adenylyl cyclases (ACs) and phosphodiesterases (PDEs) [25]. ACs, which synthesize cAMP from ATP, are either directly or indirectly regulated by Ca^2+^ signaling. Different AC isoforms and their spatial organization in cells are thought to play an important role in regulating the spatiotemporal dynamics of cAMP [26]. AC1 is an isoform of AC that is highly expressed in the brain and is directly activated by Ca^2+^ through the calcium-calmodulin complex. Phosphodiesterases hydrolyze cAMP to AMP and are regulated by cAMP-dependent protein kinase-A (PKA) activity downstream of cAMP [27, 28]. The isoforms PDE1 and PDE4 are highly expressed in the hippocampus and are modulated by the calcium-calmodulin complex and phosphorylation by PKA respectively [29].

The primary effector of cAMP, PKA, is a heterotetrameric protein kinase consisting of two catalytic and two regulatory subunits. Postsynaptically, calcium influx through N-methyl-D-aspartate receptors (NMDAR) triggers the synthesis of cAMP by ACs and induces PKA activity. PKA activation plays an important role in long-term potentiation and depression [30] and neurite outgrowth [31], resulting in memory formation [32] and emotional expression, among other cognitive processes [33]. Within dendritic spines, PKA is enriched following stimulation of upstream signaling pathways [34, 35] and structural plasticity [36]. Spinal PKA activity has also been associated with maintenance of synaptic strength [35]. Several mechanistic models have been developed to model cAMP signaling pathway dynamics [37–44]. These models have investigated the mechanisms by which molecular components such as NMDAR and *β*AR are involved in LTP through cAMP pathway [43,44]. Some others have studied localized cAMP signaling by PKA-mediated phosphorylation of PDE4 [40] or localization and colocalization of PKA by AKAPs in different subcellular domains (spine vs dendrite) [41, 42]. However, despite the critical role of PKA in signal transduction and physiological response within dendritic spines, little is known about how calcium-cAMP/PKA crosstalk occurs in dendritic spines, particularly in response to periodic oscillations of calcium.

One of the interesting dynamical features of calcium influx into postsynaptic spines is the frequency of oscillations [45–47]. Intracellular calcium oscillations ranging from seconds to minutes are known to be responsible for distinct physiological roles in the nervous system [48–51]. Oscillatory gating dynamics of plasma membrane-embedded channels or cyclic calcium release from intracellular stores can generate calcium spikes [23]. Changes in the amplitude and frequency of these oscillating signals along with the intracellular concentrations of these second messengers can trigger diverse cellular responses in neurons [23,52–54]. Studies have shown that calcium oscillates spontaneously in spines [55] and that CaMKII activity is sensitive to calcium frequency [56–60]. Given these observations, we asked, what, if any, are the relationships between Ca^2+^ frequency and cAMP/PKA dynamics? To answer this question, we built a mathematical model of calcium-induced cAMP/PKA dynamics that integrates key steps in the modulation of signaling components in the cAMP/PKA pathway. This model was constrained using experimental data available in the literature. Experimental measurements of calcium oscillations in dendritic spines show that calcium oscillates spontaneously in spines and these measurements are used to generate predictions of cAMP/PKA kinetics for different calcium inputs. Our model predicts that components in this pathway serve as frequency modulators of calcium dynamics such that cAMP/PKA only picks up the lower frequency of calcium influx; that is, cAMP/PKA is a leaky integrator of calcium dynamics. This finding sheds light on the kinetic mechanisms for robustness of cAMP/PKA responses in dendritic spines.

## Materials and Methods

### Experimental methods

#### Plasmids

GCaMP6f was purchased from Addgene.

#### Ethical Approval

All animal experiments in this study were conducted in accordance with the National Institutes of Health Guide for the care and use of laboratory animals and all protocols were approved by the University of California San Diego Institutional Animal Care and Use Committee. All surgical procedures were terminal and isoflurane was used as an anesthetic to prevent any pain or discomfort.

#### Rat primary hippocampal neuron culture

Hippocampi were dissected from E19 Sprague-Dawley male and female rat pups (Charles River) in ice-cold HEPES-buffered saline solution (Gibco), supplemented with pen-strep (Lifetech), glucose, and additional HEPES. Hippocampi were dissociated using papain (Worthington Biochemical). Cells were washed and resuspended in phenol red-free Neurobasal Medium (Lifetech) supplemented with L-glutamine (Stem Cell Technologies) and SM1 (NeuroCult, Stem Cell Technologies). Mixed neuron and astrocyte cell suspension was plated at a concentration of 200,000 cells/mL onto glass bottom 35 mm petri dishes (CellVis) coated with poly-L-lysine (Sigma). Cells were incubated in a humidified 37 *°*C, CO_2_ incubator (HeraCell). At DIV 20, cells were transfected with GCaMP6f using Lipofectamine 2000 (Invitrogen) for 48 hours prior to imaging.

#### Live-cell imaging

Neurons were imaged in standard medium using a resonant-scanning A1R confocal microscope (Nikon) equipped with a 60x 1.49 NA oil immersion objective, 489.1 nm laser, 525/50 nm emission filter, 488 dichroic, and DU4 GaASP detectors. Cells were kept at 37 *°*C and 5% CO_2_ using a stage-top incubation system (Okolab). Images of basal calcium dynamics were acquired using NIS Elements AR 5.10.01 at 4 frames per second for 4-5 minutes. In all experiments, images were acquired using identical parameters and settings (laser excitation power, acquisition time, time-lapse interval). We replicated the findings using at least 3 independent culture preparations.

#### Identification of dendritic spines

Dendritic spines were selected from proximal and distal regions of dendrites. The images shown in Figure 6 are portions of a dendrite image. For analysis, we selected only dendritic spines in which the head and neck, attached to the dendrite shaft, were visible in a single plane of focus, which was necessary to achieve sufficiently rapid image acquisition. Only experiments in which focus was perfectly maintained throughout the recording session were included in the time-lapse analyses. Dendritic spine selection was designed to minimize sampling bias by including approximately equal numbers of dendritic spines of different morphology, which was assessed on multiple planes of focus prior to starting the single plane acquisition.

**Figure 1:**
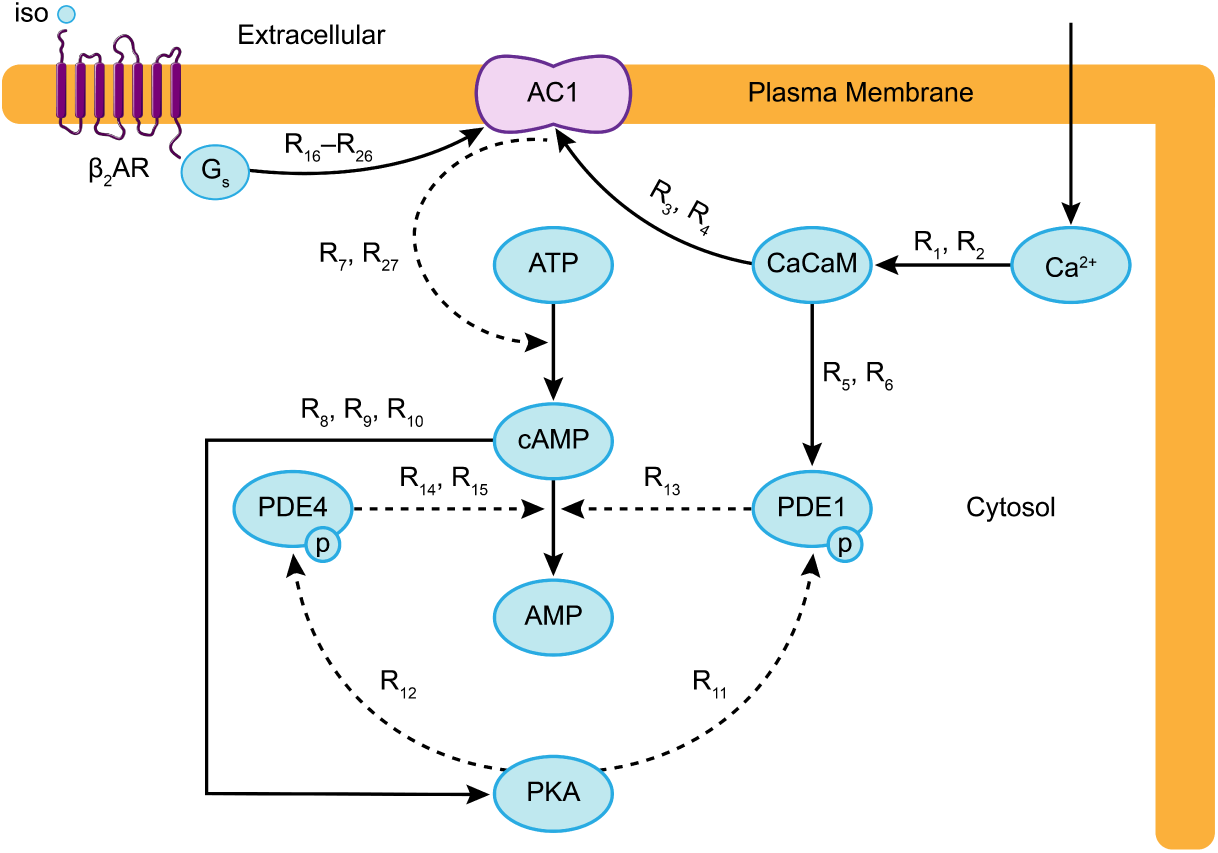
Schematic of the neuronal cAMP-PKA pathway. The main biochemical components in the pathway modeled are shown. Each arrow depicts the corresponding reactions between the different biochemical components of the model. Dashed arrows represent enzymatic reactions modeled using Michaelis-Menten kinetics and solid arrows represent reactions modeled using mass action kinetics. The reaction numbers on each arrow correspond to the reactions listed in Table 1.

**Figure 2:**
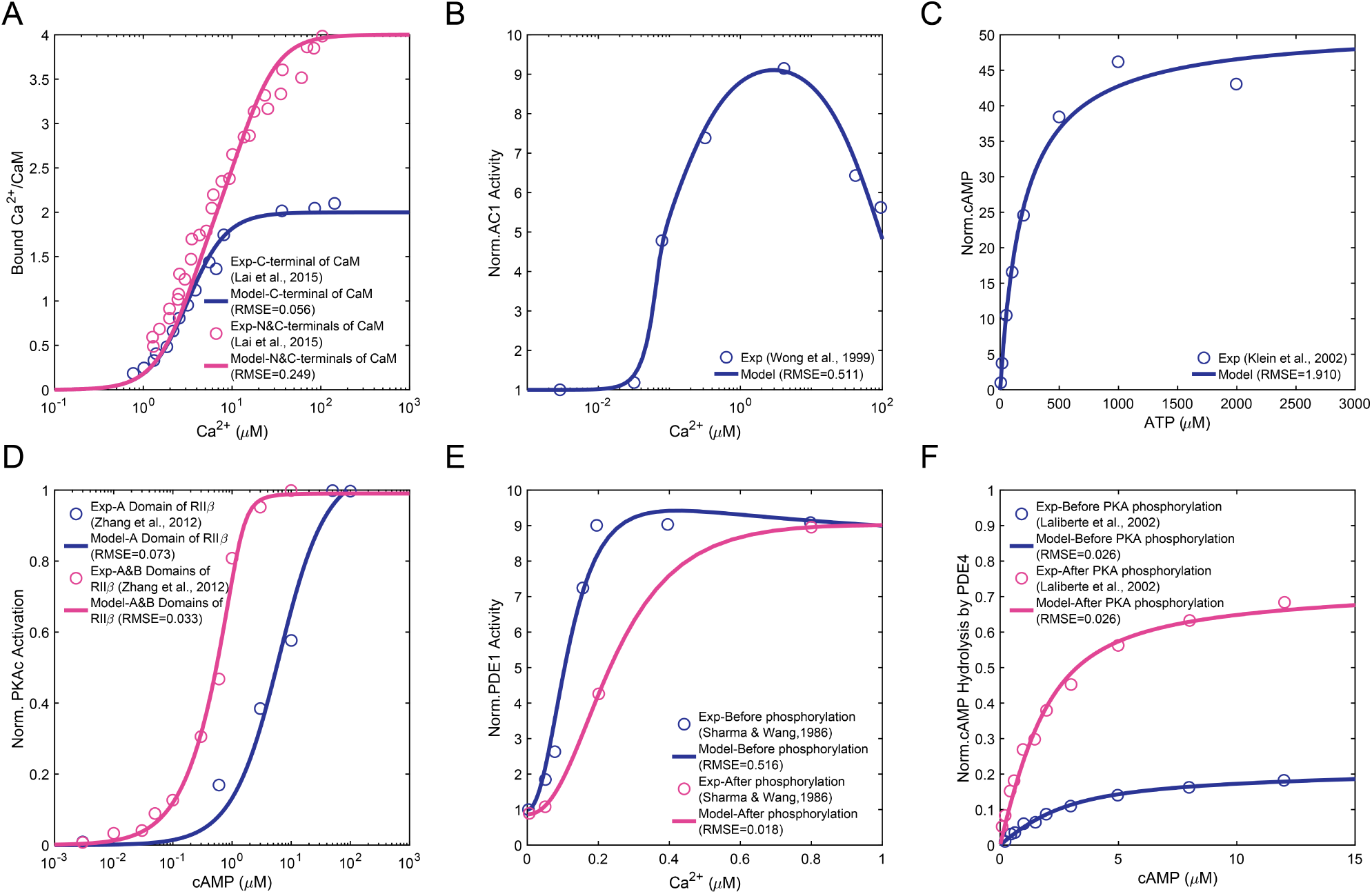
Steady-state parameter scans to evaluate the model based on available experimental data in the literature: Saturation curve of C- and N-lobes of calmodulin, predicted by our model, and comparison with experimental data from [80]. (B) Ca^2+^-stimulated adenylyl cyclase activity, predicted by our model, and comparison with experimental data from [67]. (C) cAMP production at increasing concentrations of ATP, predicted by our model, and comparison with experimental data from [68]. (D) Activation of RII*β* holoenzymes by cAMP in A- and B-domains, predicted by our model, and comparison with experimental data from [102]. (E) Activation of phosphodiesterase isoenzyme PDE1A by different concentrations of Ca^2+^ before and after the phosphorylation of the isoenzyme by PKA, predicted by our model, and comparison with experimental data from [69]. (F) PDE4A-catalyzed cAMP hydrolysis at increasing cAMP concentration before and after PKA phosphorylation, predicted by our model, and comparison with experimental data from [71]. It must be noted that none of the experimental datasets in the literature reported error bars or standard deviations for the measured data points.

**Figure 3:**
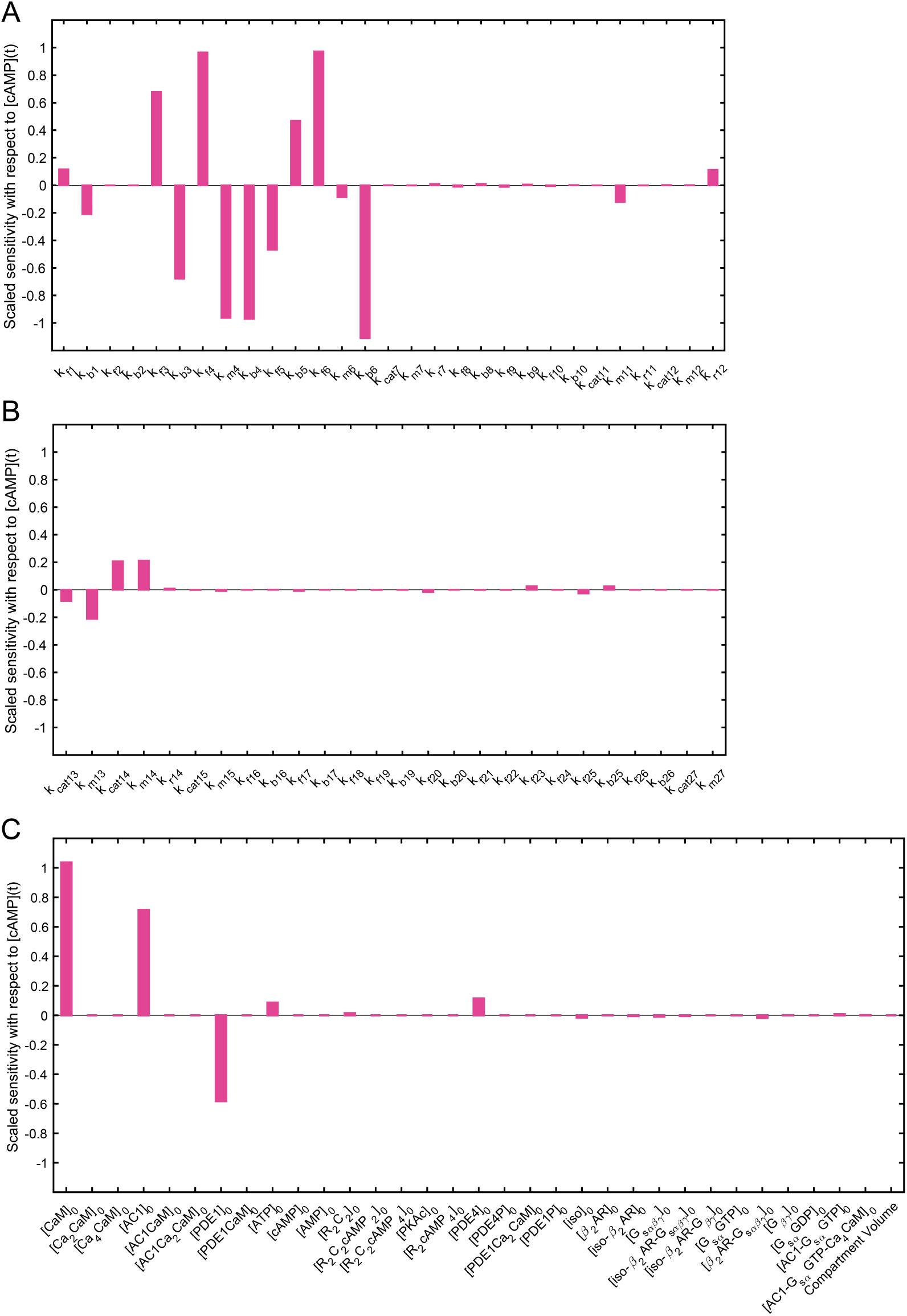
Sensitivity analysis with respect to cAMP transient concentration for (A) parameters of reactions 1 to 12, (B) parameters of reactions 13 to 27, and (C) initial concentrations and compartment volume. Kinetic parameters of reactions 3 to 6, and initial concentrations of calmodulin, AC1, and PDE1 show the highest impact on cAMP transient concentration.

**Figure 4:**
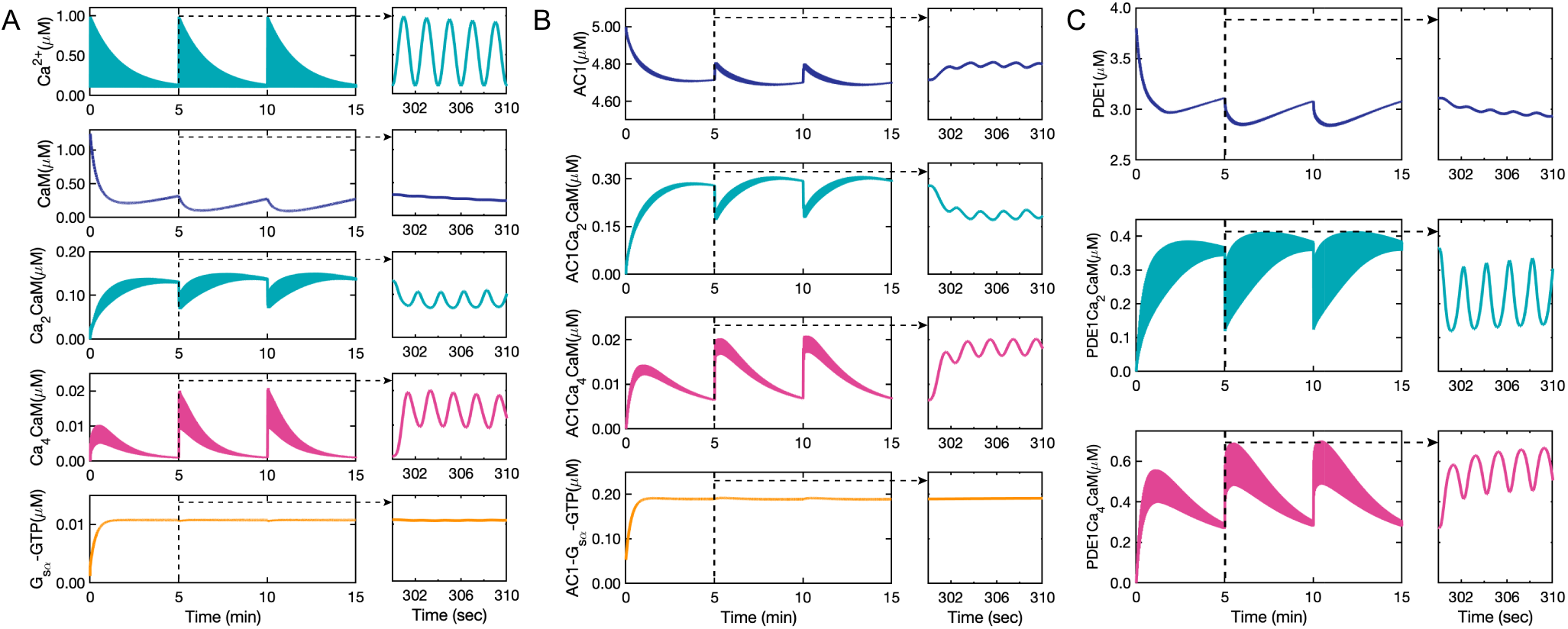
Ca^2+^/CaM complex formation and Ca^2+^/CaM-activated enzymes time course predicted by our model: (A) Calcium input with sinusoidal oscillations (0.5 Hz) and exponential decay (0.003 Hz) as the stimulus in the model. Free calmodulin concentration over a 15-minute and a 10-second time course showing only minute-scale oscillations. The first step of calcium-calmodulin binding of two calcium ions to C-lobe of calmodulin over a 15-minute and a 10-second time course. Ca^2+^ CaM shows both minute-scale (0.003 Hz) and second-scale (0.5 Hz) oscillations. The second step of calcium-calmodulin binding of another two calcium ions to N-lobe of calmodulin over a 15-minute and a 10-second time course. Ca_4_·CaM also shows both minute-scale (0.003 Hz) and second-scale (0.5 Hz) oscillations. However, G_*s*_ as the other component responsible for AC1 activation shows no oscillations as it is not activated by Ca^2+^, (B) AC1 partial activation by Ca_2_·CaM to from AC1·Ca_2_·CaM followed by binding of two extra calcium ions to fully activate AC1 and form AC1·Ca_4_·CaM. Unlike AC1 activated by Ca^2+^, AC1 activated by G_*s*_ shows no oscillations, (C) PDE1 partial activation by Ca_2_·CaM initially followed by further binding of two calcium ions to fully activate PDE1 and form PDE1·Ca_4_·CaM.

**Figure 5:**
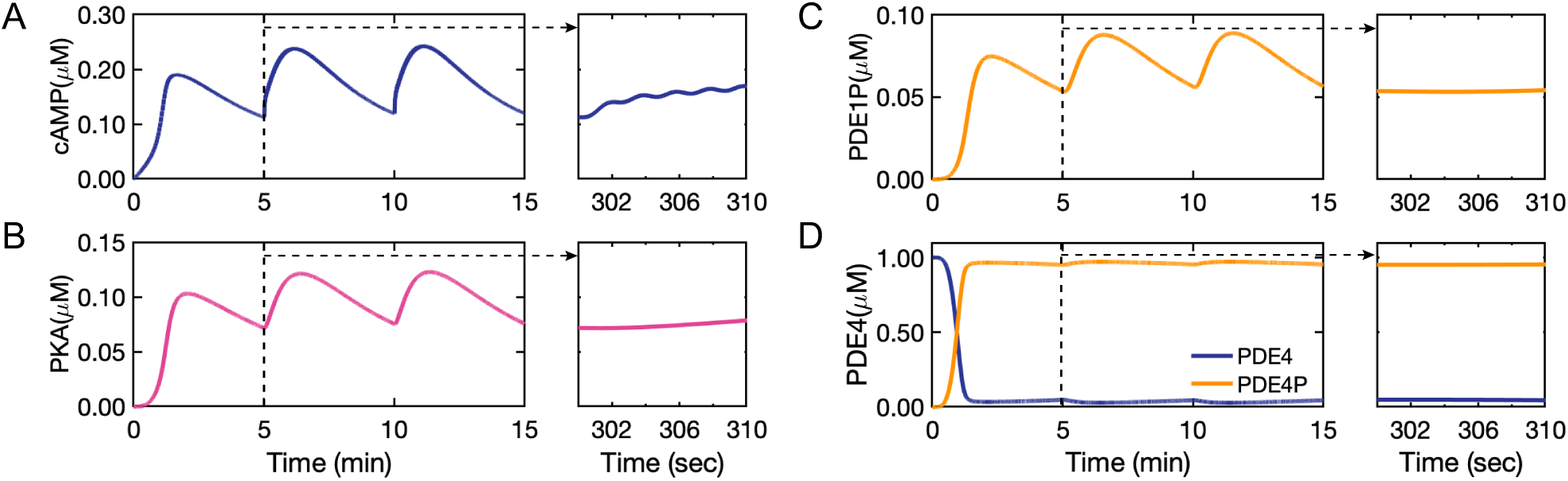
cAMP and PKA formation time course and PDE1 and PDE4 phosphorylation by PKA predicted by our model: (A) cAMP dynamics predicted by our model show mainly minute-scale oscillations leaky oscillations on the second-scale shown in the figure inset (B) PKA dynamics predicted by our model shows only minute-scale oscillations and no second-scale oscillations (figure inset). (C) Phosphorylation of PDE1 by PKA and formation of phosphorylated PDE1 (PDE1P) that oscillates only on minute-scale. (D) PDE4 and phosphorylated PDE4 (PDE4P) formed by PKA that is indirectly affected by calcium and shows almost neither of the second- and minute-scale oscillations.

**Figure 6:**
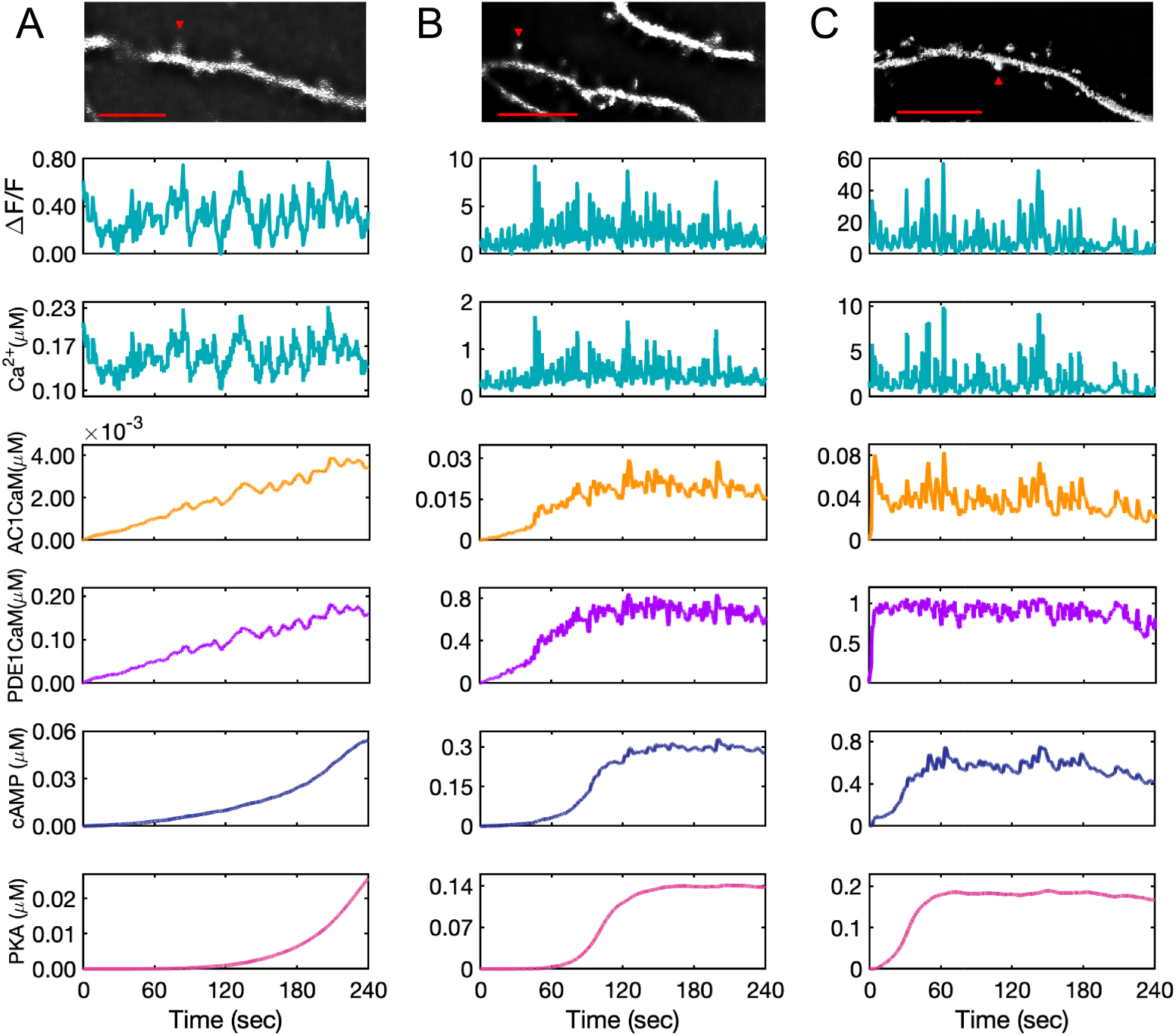
Model predictions for AC1 · CaM, PDE1 · CaM, cAMP, and PKA for experimental calcium input measured from GCaMP6f expressed in DIV 22 primary rat hippocampal neurons. Fluorescence intensity measurements shown for calcium have an arbitrary starting time and the model system at time zero (start of the recording) is already in equilibrium. The calcium concentration is estimated based on the relative increases and decreases in calcium to the minimum intensity measured. (A) The calcium input is at low range, almost on the range of resting calcium concentration. This calcium amount slows down the rate of AC1 and PDE1 activation by calcium-calmodulin complex and as a result, only a low amount of cAMP is produced. AC1 · CaM and PDE1 · CaM seem to pick up only calcium peaks, while these calcium dynamics seem to barely affect the cAMP and PKA dynamics over the time course of recorded calcium (4 minutes). (B) The second set of measured calcium in the spine shows a higher amount of calcium which accelerates the rate of AC1 and PDE1 activation and cAMP and PKA formation. Due to the higher range of calcium oscillations in this data set, AC1 · CaM, PDE1 · CaM, and cAMP seem to be affected more by calcium transients. (C) This set of measured calcium data shows the highest recorded calcium. Because of the high range of calcium oscillations (10 *µM*), the induced calcium transients are significant in this case and even PKA shows a subtle effect of the most prominent calcium spikes. The data are representative of n=10 neurons from three independent experiments. Representative images at the indicated time points are shown for each spine selected for analysis. Scale bar, 10 *µm*.

#### Image Analysis

The goal of the live cell imaging data was to verify the existence of calcium oscillations in post-synaptic spines and to extract the frequency of calcium oscillations as input for the model to test model robustness. Initial image analysis was done using NIS Elements AR Analysis 5.10.01. To measure calcium in spines, images were deconvoluted in 2D using 20 iterations of blind deconvolution with medium noise. Individual spines were identified and intensity measured over time. Fluorescence intensities were background corrected by subtracting intensity from a region with no transfected cells from spine intensities and represented as Δ*F/F* where the minimum fluorescence intensity was subtracted from the instantaneous fluorescence intensity and divided by the minimum fluorescence intensity. Further processing was done using Microsoft Excel and MATLAB. Figure preparation was done using OMERO.

### Mathematical Model Development

In order to simulate the temporal dynamics of calcium-stimulated cAMP/PKA pathway in dendritic spines, we developed a well-mixed model that accounts for the dynamics of the different biochemical species shown in Figure 1. Here, we list the model assumptions, identification of the key components in the pathway, and the governing equations.

### Model assumptions

- **Compartment size:** We model the spine as a single compartment with a volume of 0.1 femtoliter (10^*−*16^ liter) based on the average volume of a single dendritic spine [61]. All the biochemical species including the membrane-bound ones are well-mixed across this compartment. A separate spatial model is presented in [62].
- **Time scales:** We focus on cAMP/PKA dynamics at the minute timescale. Calcium input was prescribed as sinusoidal functions with exponential decays based on experimental observations in the literature [63] and our experimental measurements. Different calcium inputs are described in detail below.
- **Reaction types:** In general, the catalytic reactions such as enzymatic reactions, phosphorylation by protein kinases, and dephosphorylation by phosphatases are modeled using Michaelis-Menten kinetics. All other reactions are modeled using mass action kinetics assuming that the concentrations are present in large quantities. The binding reactions are modeled using either mass action or cooperative kinetics. Catalytic reactions representing enzymatic activation or inhibition are modeled using Michaelis-Menten kinetics. For binding/unbinding reactions we started with mass-action kinetics and then if experimental data showed high nonlinearity or cooperativity, we used kinetic rates with higher nonlinearities (Figure S1). For enzyme activations, especially the final step of activations we used Michaelis-Menten kinetics. The reactions and corresponding reaction rates are given in Table 1.
- **Enzyme regulations by calcium-calmodulin complex:** We assumed that the enzymes regulated by calcium-calmodulin, AC1 and PDE1, are fully activated only when all four binding sites of calmodulin are bound by calcium ions [64, 65]. These fully activated enzymes are shown as AC1·CaM (AC1 · Ca_4_·CaM) and PDE1·CaM (PDE1·Ca_4_·CaM).
- **Kinetic parameters:** Kinetic parameters for these reactions were estimated using the COPASI simulator [66] and are given in Table 2. The model was constrained such that dose-response curves from the simulations matched the experimentally reported dose-response curves. To estimate the model parameters, we primarily used data from brain tissues. For AC1 activation by Ca^2+^ we used mouse hippocampus data [67]. For cAMP-ATP dynamics data from mouse neuroblastoma (N18TG2 cells) was used [68]. For PDE1 activation and PDE1/PKA phosphorylation, we used data from bovine brain (60 kDa) [69,70]. In cases where data from brain tissues was not available, we used other cell types. For example, for PDE4/PKA phosphorylation, we used data from SF9 cells (spodoptera frugiperda Sf21 insect cells) [71]. The vast majority of experimental data available in the literature is based on dose-response curves and steady-state data. We mostly found steady-state data to fit the parameters since time course data was not often available in the literature. Although kinetic parameters have not been fitted to time course data, the oscillation time scales and concentration levels have been considered by constraining the lower bound and upper bound of calculated parameters.

**Table 2:** Reaction parameters calculated for the model

| Reaction rate | $k_f$ | unit | $k_b$ | unit | $K_{cat}$ | unit | $K_m$ | unit | $K_r$ | unit |
| --- | --- | --- | --- | --- | --- | --- | --- | --- | --- | --- |
| $R_1$ | 0.1 | $\mu M^{-2} \cdot s^{-1}$ | 1 | $s^{-1}$ | - | - | - | - | - | - |
| $R_2$ | 0.005 | $\mu M^{-2} \cdot s^{-1}$ | 1 | $s^{-1}$ | - | - | - | - | - | - |
| $R_3$ | 45.46 | $\mu M^{-1} \cdot s^{-1}$ | 1 | - | - | - | - | - | - | - |
| $R_4$ | 6.89 | $s^{-1}$ | 0.39 | $s^{-1}$ | - | - | 93.02 | $\mu M$ | - | - |
| $R_5$ | 87.65 | $\mu M^{-1} \cdot s^{-1}$ | 1 | $s^{-1}$ | - | - | - | - | - | - |
| $R_6$ | 49.80 | $s^{-1}$ | 1.09 | $s^{-1}$ | - | - | 6.95 | $\mu M$ | - | - |
| $R_7$ | - | - | - | - | 7.41 | $s^{-1}$ | 193.73 | $\mu M$ | 0.44 | $s^{-1}$ |
| $R_8$ | 0.19 | $\mu M^{-2} s^{-1}$ | 1 | $s^{-1}$ | - | - | - | - | - | - |
| $R_9$ | 27.89 | $\mu M^{-2} s^{-1}$ | 1 | $s^{-1}$ | - | - | - | - | - | - |
| $R_{10}$ | 0.21 | $s^{-1}$ | 1 | $\mu M^{-2} s^{-1}$ | - | - | - | - | - | - |
| $R_{11}$ | - | - | - | - | 0.10 | $s^{-1}$ | 0.42 | $\mu M$ | 0.12 | $s^{-1}$ |
| $R_{12}$ | - | - | - | - | 7.67 | $s^{-1}$ | 1.34 | $\mu M$ | 0.02 | $s^{-1}$ |
| $R_{13}$ | - | - | - | - | 1* | $s^{-1}$ | 35* | $\mu M$ | 0 | $s^{-1}$ |
| $R_{14}$ | - | - | - | - | 8.66 | $s^{-1}$ | 1.21 | $\mu M$ | 0.10 | $s^{-1}$ |
| $R_{15}$ | - | - | - | - | 1.59 | $s^{-1}$ | 0.84 | $\mu M$ | 0 | $s^{-1}$ |
| $R_{16}$ | 5.556 | $\mu M^{-1} \cdot s^{-1}$ | 5 | $s^{-1}$ | - | - | - | - | - | - |
| $R_{17}$ | 0.6 | $\mu M^{-1} \cdot s^{-1}$ | 0.001 | $s^{-1}$ | - | - | - | - | - | - |
| $R_{18}$ | 20 | $s^{-1}$ | - | - | - | - | - | - | - | - |
| $R_{19}$ | 0.04 | $\mu M^{-1} \cdot s^{-1}$ | 0.0003 | $s^{-1}$ | - | - | - | - | - | - |
| $R_{20}$ | 2.5 | $\mu M^{-1} \cdot s^{-1}$ | 0.5 | $s^{-1}$ | - | - | - | - | - | - |
| $R_{21}$ | 20 | $s^{-1}$ | - | - | - | - | - | - | - | - |
| $R_{22}$ | 80 | $s^{-1}$ | - | - | - | - | - | - | - | - |
| $R_{23}$ | 1 | $s^{-1}$ | - | - | - | - | - | - | - | - |
| $R_{24}$ | 100 | $\mu M^{-1} \cdot s^{-1}$ | - | - | - | - | - | - | - | - |
| $R_{25}$ | 38.5 | $\mu M^{-1} \cdot s^{-1}$ | 10 | $s^{-1}$ | - | - | - | - | - | - |
| $R_{26}$ | 6 | $\mu M^{-1} \cdot s^{-1}$ | 0.9 | $s^{-1}$ | - | - | - | - | - | - |
| $R_{27}$ | 10 | $\mu M^{-1} \cdot s^{-1}$ | 2273 | $s^{-1}$ | 56.84 | $s^{-1}$ | 233† | $\mu M$ | - | - |

\*All the kinetic parameters are calculated by fitting the experimental data to the kinetic equations. The kinetic parameters of reaction 13 are directly extracted from [70] and the kinetic parameters of reactions 16-27 are directly extracted from [43].

Based on these assumptions, we constructed a well-mixed model of the calcium-induced cAMP/PKA pathway in dendritic spines.

### Key components and governing equations of the model

The primary focus of this study is the regulation of cAMP by predominant Ca^2+^-sensitive isoforms of ACs and PDEs in the brain and more specifically in the hippocampus. The isoforms of adenylyl cyclase expressed at high levels in rat brain are AC1, AC2, and AC5 [72]. AC1 is predominantly found in hippocampus and cerebellum, AC5 is restricted to the basal ganglia, and AC2 is more widely expressed, but at much lower levels [72]. The predominantly Ca^2+^/calmodulin-stimulated AC1 is selectively expressed in areas associated with learning and memory, including cortex, hippocampus, and cerebellum. The Ca^2+^-inhibitable AC5 is exclusively expressed in striatal medium-sized neurons [72]. Therefore, only AC1 has been considered in this model. Of all the eleven known members of the PDE family, only PDE1 and PDE4 are directly regulated by a rise in calcium [73]. After identifying the key components of the cAMP pathway, using the existing experimental data in the literature, we designed the the model using the following eight modules:

1. **Ca**^2+^**/CaM complex formation:** Calcium can enter the spine through NMDAR-mediated influx at the plasma membrane or through the activation of ryanodine and IP_3_ receptors from internal stores [74]. Resting calcium is in the range of 0.05-0.1 *µ*M and can rise to 10-100-fold when neurons are activated [75]. Calmodulin is a small protein involved in the regulation of many calcium-dependent events [76–78]. Calmodulin has two calcium binding sites at each N-terminus and C-terminus lobe with different Ca^2+^ affinities [79]. Calcium has a higher affinity for C-lobes than the N-lobes [80] but the N-lobe binds calcium faster [81]. Based on these observations, we modeled the formation of the calcium-calmodulin complex using two sequential binding reactions with mass action kinetics. For simplicity, we did not consider the cooperative binding within each of these individual lobes [82]. Reaction 1 in Table 1 accounts for binding of the first two calcium ions to the C-lobe resulting in the formation of Ca_2_ · CaM and reaction 2 in Table 1 accounts for the binding of third and fourth calcium ions to the N-lobe resulting in the formation of Ca_4_ · CaM. Kinetic parameters of these reactions (kf_1_, kb_1_, kf_2_, kb_2_) were obtained by fitting experimental data [80] to the model in COPASI (Table 2, Figure 2A).
2. **AC1 activation by Ca**^2+^**/CaM complex:** Calcium-calmodulin stimulated adenylyl cyclase 1 (AC1) plays an important role in learning, memory formation, and long-term potentiation by coupling calcium activity to cAMP in neurons [83]. Unlike other members of AC family, AC1 is not stimulated by G_*s*_-coupled receptors unless it is activated by intracellular calcium [84]. AC1 responds to NMDA-mediated calcium elevations (and not calcium released from intracellular stores) and is mostly located close to the post-synaptic densities of dendritic spines [85]. AC1 is the predominant and neuro-specific calcium-stimulated adenylyl cyclase which is mostly expressed in dendrites, axons and at synapses [86]. At very high concentrations of calcium (*>* 10 *µM*) AC1 is inhibited [87]. Our model takes this effect into account by fitting experimental data from [67] to the AC1 activity for steady state Ca^2+^ (Figure 2B). AC1 synthesizes cAMP from ATP and has been shown to be associated with hippocampal-dependent learning abilities [85]. We modeled AC1 activation by calcium as a two-step process. First, AC1 binds to Ca_2_ · CaM by mass action kinetics to form AC1 · Ca_2_ · CaM (reaction 3 in Table 1). Second, two other Ca^2+^ bind to AC1 · Ca_2_ · CaM in a cooperative binding reaction to form AC1 · Ca_4_ · CaM (reaction 4 in Table 1). Only fully activated AC1 · Ca_4_ · CaM can catalyze cAMP production. The first term in R_4_ accounts for cooperative kinetics, while the second term accounts for basal degradation. As previously mentioned, the calcium/calmodulin binding in this study was simplified to two binding reactions: the first two calcium binding calmodulin and producing Ca_2_ · CaM and then Ca_2_ · CaM binding to two more calcium ions and producing Ca_4_ · CaM. Based on Masada *et al.* [64] study showing AC1 is not activated by half-occupied calmodulin, we assumed that only the product of the second reaction (Ca_4_ · CaM) can activate AC1. The parameters used in this module are listed in Table 2 and comparison against experimental data is shown in Figure 2B.
3. **PDE1 activation by Ca**^2+^**/CaM complex:** Phosphodiesterase 1 (PDE1) is the other key enzyme that regulates calcium/cAMP interactions [65]. PDE1A (the PDE1 subtype used in our model) is widely expressed at high levels in the hippocampus [88]. PDE1 is the only type of phosphodiesterase that is activated by calcium-calmodulin [89, 90]. PDE1 becomes activated by an increase in intracellular calcium level and has low cAMP affinity in comparison with other members of phosphodiesterase family [91]. PDE1 isozymes show low cAMP affinities and low basal PDE1 activities. Based on the study by Sharma *et al.* [92] the isozymes function mainly during cell activations when both cAMP and Ca^2+^ concentrations are elevated. The phosphorylation of PDE1A (brain 60 kDa PDE1) by PKA increases the calcium concentration required for activation of PDE1A by decreasing the enzyme’s affinity for calmodulin [92]. Unlike AC1, PDE1A does not discriminate between different sources of calcium [73]. We modeled PDE1 activation by calcium as a two-step process – first, PDE1 binds to Ca_2_ · CaM by mass action kinetics to form PDE1 · Ca_2_ · CaM (reaction 5 in Table 1). Second, two more Ca^2+^ bind to PDE1 · Ca_2_ · CaM in a cooperative binding reaction to form PDE1 · Ca_4_ · CaM (reaction 6 in Table 1). Only fully activated PDE1 · Ca_4_ · CaM can hydrolyze cAMP into AMP. The first term in R_6_ accounts for cooperative kinetics, while the second term accounts for basal degradation. The parameters used in this module are listed in Table 2 and comparison against experimental data from [69] is shown in Figure 2E.
4. **cAMP production:** Adenylyl cyclases catalyze the conversion of ATP into cAMP. cAMP is implicated in both LTP and LTD by activating PKA [93–96]. In our model, fully activated AC1, AC1 · Ca_4_ · CaM, catalyzes the formation of cAMP from ATP (reaction 7 in Table 1). This reaction is modeled by Michaelis-Menten kinetics with a basal degradation term. The basal activity of the type I adenylyl cyclase (AC1) is regulated by calcium/calmodulin [97]. In the absence of Ca^2+^ influx, AC1 shows a low level of basal activity, producing modest levels of cAMP [98]. Therefore, the AC1 basal cAMP production has not been explicitly modeled as an individual reaction. The steady-state amount of generated cAMP with ATP is evaluated using experimental data from [68] shown in Figure 2C.
5. **PKA activation:** PKA has two catalytic subunits and a regulatory dimer that binds to the catalytic subunits [99, 100]. RII*β* is the dominant PKA isoform expressed in the brain [101]. cAMP binds to its effector, PKA, and regulates the activity of regulatory subunits. Then, PKA phosphorylates different protein kinases responsible for neuron growth and differentiation, such as LKB1 and GSK-3*β* in the hippocampus [91]. Activation of PKA by cAMP is modeled in three steps inspired by [101] with some modifications in the order of A- and B-domain binding of cAMP that was observed in experiments by [102]. First, two cAMP molecules bind to the A-domain (reaction 8 in Table 1), then the other two cAMP molecules bind to the B-domain (reaction 9 in Table 1). Finally, when all four cAMP molecules are bound to A- and B-domains, the two catalytic subunits are released (reaction 10 in Table 1). These reaction schemas are evaluated by experimental data from [102] for steady state cAMP concentration and PKA formation shown in Figure 2D and kinetic parameters are given in Table 2.
6. **PDE phosphorylation by PKA:** Both PDE1A and PDE4A can be phosphorylated by PKA. PKA phosphorylation of PDE1A alters its affinity toward both calcium and calmodulin [69]. PDE4 is a cAMP-specific phosphodiesterase and its activity is increased two to six-fold upon PKA phosphorylation [91]. Different PDE4 isoforms (PDE4A, PDE4B, PDE4D) are targeted to synapses to regulate the cAMP level and modulate synaptic plasticity in learning and memory [91]. PDE4A (the PDE4 subtype used in our model) is highly expressed in the CA1 subregions of the rat hippocampus and is the major PDE4 subtype involved in mediating memory [88]. The phosphorylation reactions of the two different PDE isoforms are modeled by Michaelis-Menten kinetics with a basal degradation, where PKA acts as an enzyme in both of these reactions (reactions 11 and 12 in Table 1). These reactions are evaluated at steady state to show how PKA phosphorylation affects the activity of PDE1A and PDE4 in comparison with experiments from [69] and [71] in Figure 2E and Figure 2F respectively.
7. **cAMP degradation:** We assumed that cAMP can be degraded to AMP by PDE1 · Ca_4_ · CaM, PDE4, and PDE4P (PDE4 phosphorylated by PKA). All cAMP inhibition reactions are modeled by Michaelis-Menten kinetics with a basal degradation term, in which phosphodiesterases act as enzymes (reactions 13-15 in Table 1). cAMP hydrolysis by PDE4 and PDE4P evaluated against experimental data from [71] is shown in Figure 2F.
8. **AC1 activation by G**_*s*_: There is some evidence indicating that the activity of NMDAR may be potentiated by stimulating GPCRs. Raman at al. [103] have shown that stimulation of *β*-adrenergic receptors during excitatory synaptic transmission can increase charge transfer and Ca^2+^ influx through NMDA receptors. Gereau and Conn [104] have also tested the hypothesis that coactivation of mGluRs and a G_*s*_-coupled receptor (the beta-adrenergic receptor) could lead to large increases in cAMP accumulation in hippocampus and thereby increasing synaptic responses in area CA1. They report that coactivation of mGluRs and beta-adrenergic receptors is not accompanied by an increase in excitatory postsynaptic currents or by a decrease in synaptic inhibition in area CA1, suggesting that it is not mediated by a lasting change in excitatory or inhibitory synaptic transmission. However, coactivation of these receptors leads to a persistent depolarization of CA1 pyramidal cells with a concomitant increase in input resistance. Gs can have an additional effect on AC1 activation. Stimulation of *β*-adrenergic receptors (*β*ARs) by an agonist such as isoproterenol can activate the G_*s*_ subtype of GTP-binding protein, which stimulates adenylyl cyclase isoforms [105]. A study by Wayman *et al.* [84] suggests that AC1 enzyme is stimulated by Gs-coupled receptors in vivo when it is activated by intracellular calcium. Based on these experimental observations, reactions 15 to 27 (Table 1) has been reproduced from [43] exactly as is to take into account the effect of AC1 activation and cAMP production by G_*s*_.

The list of the above reactions and estimated parameters are shown in Table 1 and Table 2, respectively.

### Ordinary differential equations

The temporal dynamics of each species, *c*, is defined as

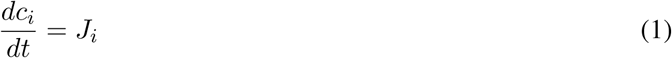

where *c*_*i*_, *i ∈* 1, 2, *…*, 21, represents the concentration of the *i*^*th*^ species as a function of time and *J*_*i*_ is the net reaction flux for the *i*^*th*^ species. These ordinary differential equations (ODEs) and the corresponding initial concentration of the different species are shown in Table 3. The sensitivity analysis with respect to kinetic parameters and initial conditions of the model with respect to transient cAMP concentration is shown in Figure 3.

**Table 3:**
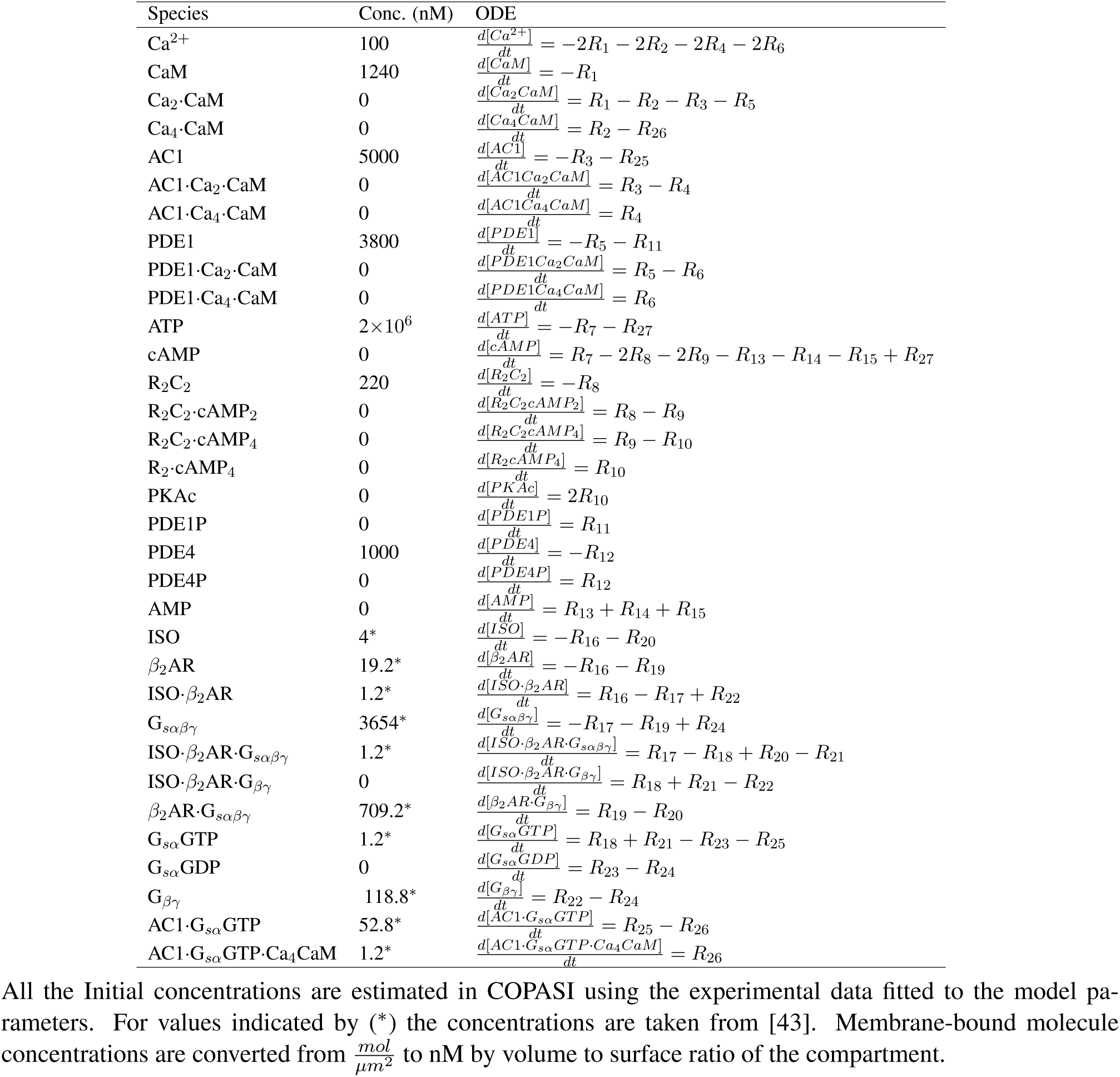
Initial concentrations and ordinary differential equations

| Species | Conc. (nM) | ODE |
| --- | --- | --- |
| Ca<sup>2+</sup> | 100 | $\frac{d[Ca^{2+}]}{dt} = -2R_1 - 2R_2 - 2R_4 - 2R_6$ |
| CaM | 1240 | $\frac{d[CaM]}{dt} = -R_1$ |
| Ca<sub>2</sub>·CaM | 0 | $\frac{d[Ca_2CaM]}{dt} = R_1 - R_2 - R_3 - R_5$ |
| Ca<sub>4</sub>·CaM | 0 | $\frac{d[Ca_4CaM]}{dt} = R_2 - R_{26}$ |
| AC1 | 5000 | $\frac{d[AC1]}{dt} = -R_3 - R_{25}$ |
| AC1·Ca<sub>2</sub>·CaM | 0 | $\frac{d[AC1Ca_2CaM]}{dt} = R_3 - R_4$ |
| AC1·Ca<sub>4</sub>·CaM | 0 | $\frac{d[AC1Ca_4CaM]}{dt} = R_4$ |
| PDE1 | 3800 | $\frac{d[PDE1]}{dt} = -R_5 - R_{11}$ |
| PDE1·Ca<sub>2</sub>·CaM | 0 | $\frac{d[PDE1Ca_2CaM]}{dt} = R_5 - R_6$ |
| PDE1·Ca<sub>4</sub>·CaM | 0 | $\frac{d[PDE1Ca_4CaM]}{dt} = R_6$ |
| ATP | 2 × 10<sup>6</sup> | $\frac{d[ATP]}{dt} = -R_7 - R_{27}$ |
| cAMP | 0 | $\frac{d[cAMP]}{dt} = R_7 - 2R_8 - 2R_9 - R_{13} - R_{14} - R_{15} + R_{27}$ |
| R<sub>2</sub>C<sub>2</sub> | 220 | $\frac{d[R_2C_2]}{dt} = -R_8$ |
| R<sub>2</sub>C<sub>2</sub>·cAMP<sub>2</sub> | 0 | $\frac{d[R_2C_2cAMP_2]}{dt} = R_8 - R_9$ |
| R<sub>2</sub>C<sub>2</sub>·cAMP<sub>4</sub> | 0 | $\frac{d[R_2C_2cAMP_4]}{dt} = R_9 - R_{10}$ |
| R<sub>2</sub>·cAMP<sub>4</sub> | 0 | $\frac{d[R_2cAMP_4]}{dt} = R_{10}$ |
| PKAc | 0 | $\frac{d[PKAc]}{dt} = 2R_{10}$ |
| PDE1P | 0 | $\frac{d[PDE1P]}{dt} = R_{11}$ |
| PDE4 | 1000 | $\frac{d[PDE4]}{dt} = -R_{12}$ |
| PDE4P | 0 | $\frac{d[PDE4P]}{dt} = R_{12}$ |
| AMP | 0 | $\frac{d[AMP]}{dt} = R_{13} + R_{14} + R_{15}$ |
| ISO | 4* | $\frac{d[ISO]}{dt} = -R_{16} - R_{20}$ |
| β<sub>2</sub>AR | 19.2* | $\frac{d[\beta_2AR]}{dt} = -R_{16} - R_{19}$ |
| ISO·β<sub>2</sub>AR | 1.2* | $\frac{d[ISO\cdot\beta_2AR]}{dt} = R_{16} - R_{17} + R_{22}$ |
| G<sub>sαβγ</sub> | 3654* | $\frac{d[G_{s\alpha\beta\gamma}]}{dt} = -R_{17} - R_{19} + R_{24}$ |
| ISO·β<sub>2</sub>AR·G<sub>sαβγ</sub> | 1.2* | $\frac{d[ISO\cdot\beta_2AR\cdot G_{s\alpha\beta\gamma}]}{dt} = R_{17} - R_{18} + R_{20} - R_{21}$ |
| ISO·β<sub>2</sub>AR·G<sub>βγ</sub> | 0 | $\frac{d[ISO\cdot\beta_2AR\cdot G_{\beta\gamma}]}{dt} = R_{18} + R_{21} - R_{22}$ |
| β<sub>2</sub>AR·G<sub>sαβγ</sub> | 709.2* | $\frac{d[\beta_2AR\cdot G_{s\alpha\beta\gamma}]}{dt} = R_{19} - R_{20}$ |
| G<sub>sα</sub>GTP | 1.2* | $\frac{d[G_{s\alpha}GTP]}{dt} = R_{18} + R_{21} - R_{23} - R_{25}$ |
| G<sub>sα</sub>GDP | 0 | $\frac{d[G_{s\alpha}GDP]}{dt} = R_{23} - R_{24}$ |
| G<sub>βγ</sub> | 118.8* | $\frac{d[G_{\beta\gamma}]}{dt} = R_{22} - R_{24}$ |
| AC1·G<sub>sα</sub>GTP | 52.8* | $\frac{d[AC1\cdot G_{s\alpha}GTP]}{dt} = R_{25} - R_{26}$ |
| AC1·G<sub>sα</sub>GTP·Ca<sub>4</sub>CaM | 1.2* | $\frac{d[AC1\cdot G_{s\alpha}GTP\cdot Ca_4CaM]}{dt} = R_{26}$ |

All the Initial concentrations are estimated in COPASI using the experimental data fitted to the model parameters. For values indicated by (\*) the concentrations are taken from [43]. Membrane-bound molecule concentrations are converted from $\frac{mol}{\mu m^2}$ to nM by volume to surface ratio of the compartment.

### Ca^2+^ input signals as the stimulus

The four different types of input calcium are modeled as:

1. Sinusoidal oscillations (*ω*_1_=0.5 Hz) with exponential decay that is repeated every 5 minutes (*ω*_2_=0.003 Hz):

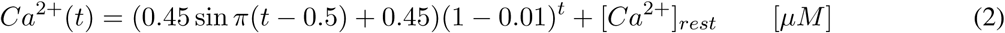
2. Calcium bursts with exponential decay that is being repeated every 5 minutes (*ω*_2_=0.003 Hz):

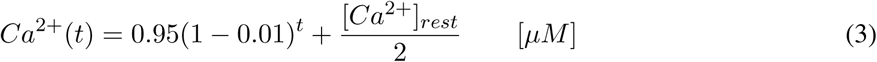
3. Non-oscillating calcium:

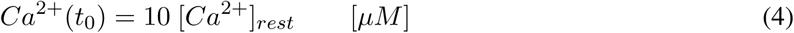

in which [Ca^2+^]_*rest*_ =0.1 *µM* is the resting concentration of calcium in the cytoplasm.
4. Experimental measurements of calcium: Spontaneous calcium oscillations measured in neurons are shown in Figure 6. The measured calcium is in the range of 0.1 to 10 *µM* with oscillations in the seconds timescale.

### Numerical Methods

The system of ordinary differential equations for this model is solved for a time course of 1800 seconds and interval size of 0.1 (sec) through a deterministic simulation in COPASI [106]. Using a deterministic algorithm (LSODA), the system of ODEs is solved with a dense or banded Jacobian when the problem is stiff and automatically selects between non-stiff (Adams) and stiff (BDF) methods by initially using the non-stiff method and dynamically monitoring data in order to decide which method to use [107]. Parameter estimations were conducted in COPASI using the Evolutionary Programming (EP) method [108–110]. The link to the model will be available on https://github.com/donya26/cAMP-PKA-temporal.git.

### Sensitivity Analysis

In sensitivity analysis of kinetic parameters with respect to transient cAMP concentration (Figure 3), kinetic parameters of reactions related to activation of AC1 and PDE1 enzymes by Ca^2+^/CaM complex (reactions 3-6) show the most significant effect on cAMP concentration (Figure 3A). However, kinetic parameters related to AC1 activation by G_*s*_ shows almost no effect on cAMP concentration (Figure 3B). Finally, initial concentration of these Ca^2+^-sensitive enzymes (AC1 and PDE1) and calmodulin shows the most significant effect on cAMP concentration (Figure 3C). It is important to note that the compartment size has a trivial effect on the model (Figure 3C). These results show the significance of regulation of cAMP/PKA pathway through Ca^2+^-sensitive enzyme activations. Sensitivity analyses were conducted in COPASI by numerical differentiation using finite differences [111].

### Stochastic Simulations

Spines are small compartments and neuronal activity in the spines is mediated through changes in the probability of stochastic transitions between open and closed states of ion channels [112]. In this model, the focus has been on cAMP regulation and for the sake of simplicity, we did not consider such stochastic calcium-related events. However, in order to make sure that the stochasticity will not change the obtained results, we also simulated the model stochastically (*τ*-leap method-using COPASI). Although in order to conduct the simulations stochastically we had to make some changes to the model, e.g. we had to make all the reactions irreversible and change some of the reaction rates, we saw almost the same trend with deterministic and stochastic methods (Figure S2). However, in order to better capture oscillation regulation of different components of the pathway, we modeled the system deterministically.

## Results

### Large frequency modulation of immediate downstream effectors of calcium

The first module of the model focused on the formation of the calcium-calmodulin complex. Calcium input was modeled as a sinusoidal function with a 2-second oscillation period with an exponential decay every 5-minutes [63] (Figure 4A). From here onwards, we refer to the larger frequency as *ω*_1_ for the oscillations in the seconds timescale and the smaller frequency as *ω*_2_ for the oscillations in the minutes timescale. The dynamics of calmodulin consumption leading to the formation of Ca_2_ · CaM respond only to the low-frequency oscillations of calcium in minute-scale (Figure 4A). Ca_2_ · CaM demonstrates both low frequency (minute-scale) and high frequency (second-scale) oscillations (Figure 4). In the next step, two more Ca^2+^ bind to Ca_2_ · CaM to form Ca_4_ · CaM. Ca_4_ · CaM also responds to second-scale oscillations of calcium; however, the oscillation amplitudes are almost 10-fold lower in comparison with Ca_2_ · CaM (Figure 4A). Another key difference between the observed oscillation patterns of Ca_2_ · CaM and Ca_4_ · CaM is that the Ca_2_ · CaM oscillation pattern seems to be mainly driven by CaM on the minute-scale, while the Ca_4_ · CaM oscillation pattern looks almost exactly like calcium patterns with less significant oscillations on the second scale.

AC1 and PDE1 are activated by the Ca^2+^/CaM complex. Ca_2_ · CaM binds to AC1 and forms AC1 · Ca_2_ · CaM. Both AC1 and AC1 · Ca_2_ · CaM show second and minute-scale calcium oscillations (Figure 4B) and their oscillation pattern seem to be mostly deriven by Ca_2_ · CaM oscillation pattern shown in Figure 4C with less significant second-scale oscillations. In the second step, AC1 · Ca_2_ · CaM fully activates AC1 by forming AC1 · Ca_4_ · CaM with the other two Ca^2+^. In comparison to AC1 · Ca_2_ · CaM, AC1 · Ca_4_·CaM shows less significant second-scale oscillations (almost 10-fold lower) and the minute-scale oscillations seems to be driven by the calcium oscillation pattern (Figure 4B). AC1 · Ca_4_ · CaM dynamics mostly follow Ca_4_ · CaM dynamics with less sensitivity to Ca^2+^. The dynamics of PDE1 activation by Ca_2_ · CaM is very similar to AC1. However, both PDE1 · Ca_2_ · CaM and PDE1 · Ca_4_ · CaM show much higher sensitivity to second-scale oscillations of Ca^2+^ (Figure 4C).

### Small frequency modulation of cAMP/PKA and PDE phosphorylation

After enzyme activation by calcium-calmodulin complex, cAMP is synthesized by AC1 · Ca_4_ · CaM and AC1 · G_*sα*_GTP · Ca_4_ · CaM from ATP. The contribution of Ca^2+^/CaM-activated AC1 in cAMP production is much more substantial in comparison to G_*s*_-activated AC1 and cAMP produced by AC1 · G_*sα*_GTP · Ca_4_ · CaM is almost negligible (Figure S3). This is in agreement with experimental observations of Baker *et al.* [113] showing that Gs-coupled receptor does not stimulate the Gs-insensitive AC1 and Halls and Cooper [26], suggesting calcium/calmodulin binding as the main activation mechanism for AC1s. Then PKA is activated by cAMP and activated PKA phosphorylates PDE1 and PDE4. Finally, cAMP is degraded by PDE1, PDE4, and PDE4P and shows minute-scale oscillations and leaky oscillations on the second-scale (Figure 5A). PKA is activated by cAMP and as a result, it only shows minute-scale oscillations (Figure 5B). While PDE1-P (phosphorylated PDE1) shows minute-scale oscillations (Figure 5C), PDE4P (phosphorylated PDE4) barely shows any oscillations (Figure 5D). Degradation of cAMP by PDE4 and PDE4P, which are indirectly affected by calcium oscillations might filter the second-scale oscillations induced by AC1 · CaM in the cAMP synthesis and change the second-scale oscillations to leaky oscillations (Figure 5A). Thus, our model predicts that phosphodiesterases may filter the large frequency (*ω*_1_) oscillations of the upstream components driven by calcium dynamics and capture only the smaller frequency (*ω*_2_).

### cAMP/PKA pathway filtering of Ca^2+^ signals

The response of cAMP/PKA to different calcium patterns raises the following questions: do spines experience vastly varying calcium patterns? Can experimentally-measured calcium patterns be used to generate predictions from our model? In order to answer these questions, we measured basal calcium fluctuations in dendritic spines of hippocampal neurons. Cultured neurons have been previously reported to form neuronal networks *in vitro* exhibiting spontaneous action potentials [114], which can be measured using reporters of neuronal activity, such as calcium sensors. We expressed the genetically encoded calcium biosensor GCaMP6f [115] in primary rat hippocampal neurons. To quantify Ca^2+^ oscillations, we assigned calcium concentrations to correspond to changes in the measured fluorescence intensity (ΔF/F), based on previously reported intracellular calcium concentrations [75]. Using the measured calcium in three spines from three neurons as the stimulus in the model, we found our model predicts different types of cAMP/PKA dynamics based on the calcium input (Figure 6). We chose these three spines to reflect the different calcium patterns tested in Figure 7. The first spine measured had minimal changes in fluorescence intensity and displays a low range of predicted calcium concentrations (0.1 to 0.2 *µM*) (Figure 6A). There seem to be no apparent calcium spikes, and the calcium concentration is barely enough to fully activate AC1 and PDE1 and show steady-state cAMP and PKA. AC1 · CaM and PDE1 · CaM integrate some calcium peaks; however, cAMP and PKA do not exhibit significant peaks. The second spine measured shows higher fluorescence intensity which is almost 10-fold higher than the first data set (Figure 6B). In the second spine, AC1, PDE1, cAMP, and PKA reach a steady state. AC1 and PDE1 seem to reflect most of the calcium major peaks, while cAMP appears to filter out some of the less significant peaks, and PKA almost shows none of these peaks. The third spine measured showed many calcium spikes and a very high range of fluorescence intensity, which we correlated to a large range of calcium concentrations (Figure 6C). The slopes at the first 60 seconds of simulation show that the activation of AC1 and PDE1, and formation of cAMP and PKA occurs much faster in the third spine (Figure 6C) in comparison to the other two (Figure 6A and B). With high ranges of calcium transients, AC1, PDE1, and cAMP seem to integrate most of the calcium peaks and even PKA has nominal peaks (Figure 6C). Thus, we find that for experimentally derived calcium transients cAMP/PKA serve to filter out the larger frequency (*ω*_1_).

**Figure 7:**
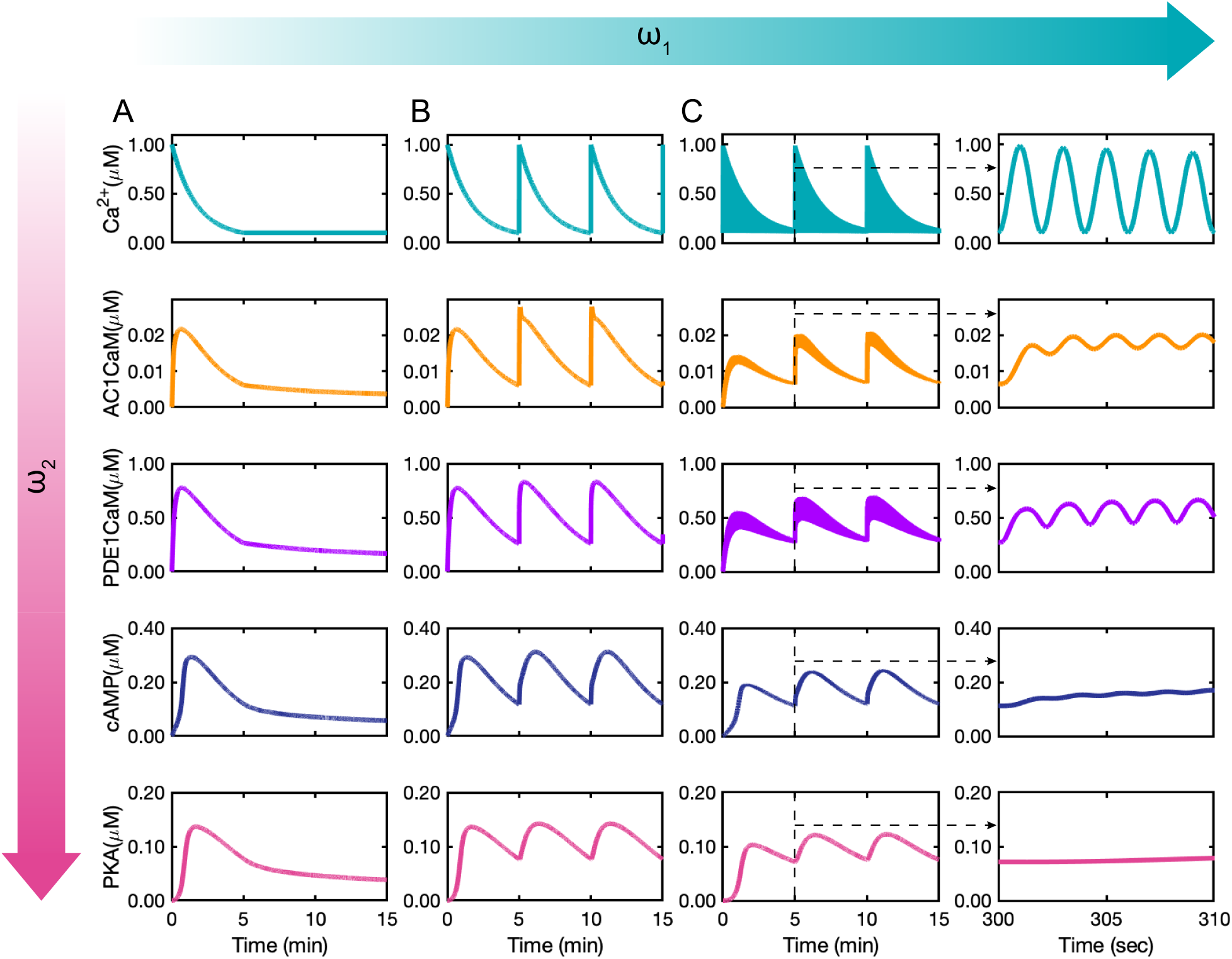
Effect of calcium stimulation patterns on AC1 · CaM, PDE1 · CaM, cAMP, and PKA dynamics: (A) Non-oscillating calcium with an initial concentration of 1 *µM* induces only one peak in each of AC1 · CaM, PDE1 · CaM, cAMP, and PKA. (B) Calcium bursts with exponential decays with 0.003 Hz frequency (*ω*_2_) induces minute-scale oscillations in each of AC1 · CaM, PDE1 · CaM, cAMP, and PKA. (C) The combination of sinusoidal calcium oscillations of 0.5 Hz with exponential decays of 0.003 Hz induces both second-scale and minute-scale oscillations in both AC1 · CaM and PDE1 · CaM as the plot insets show. While cAMP barely shows a leaky oscillation on the second-scale, PKA merely shows minute-scale oscillations.

### Ca^2+^ signal frequency modulates cAMP/PKA dynamics

If it is true that cAMP/PKA filters out *ω*_1_ dynamics to pick *ω*_2_ dynamics, then this must hold true for different values of these two frequencies. To further test our predictions, we designed three different calcium inputs (Figure 7) – first, no oscillations of calcium; second, *ω*_1_ is zero and only *ω*_2_ is present; and third, both *ω*_1_ and *ω*_2_ are present. In the first set of simulations, Ca^2+^ input has an initial constant concentration of 1 *µM* with no oscillations or calcium bursts during the 15 minutes of oscillations (Figure 7A). From AC1 · CaM and PDE1 · CaM to cAMP and PKA, it is evident that there is only one peak of the concentrations for all these species, with a 40-60 seconds delay between AC1 · CaM and PDE1 · CaM and cAMP/PKA. This behavior has also been observed by [36], where PKA returns to basal level about 10 minutes after stimulation by a pulse of calcium. For the second set of simulations, starting from the same initial calcium concentration (1 *µM*), we simulated a calcium burst every five minutes (Figure 7B). These calcium bursts at every five minutes are reflected in AC1 · CaM, PDE1 · CaM, cAMP, and PKA dynamics. The range of concentrations for all of these molecules is the same as the concentration seen in the first simulation (Figure 7A). The third set of simulations has sinusoidal oscillations with two-second periods in addition to the calcium bursts at every five minutes (Figure 7C). AC1 · CaM and PDE1 · CaM show both second- and minute-scale oscillations while cAMP and PKA only show the minute-scale oscillations (Figure 7C). In comparison to the second simulation (Figure 7B), the concentration of AC1 · CaM, PDE1 · CaM, cAMP, and PKA are lower (Figure 7C). We also chose different *ω*_2_ values and confirmed that cAMP only picks up minute time scales (Figure S4).

## Discussion

cAMP/PKA activity triggered by Ca^2+^ is an essential biochemical pathway for synaptic plasticity, regulating spine structure, and long-term potentiation [36,116]. In this study, we have developed a mathematical model, constrained by existing experimental data (Figure 2), to simulate the dynamics of cAMP/PKA pathway in response to calcium input. Analysis of this model allowed us to make the following predictions: *first*, for a given calcium input, AC1, and PDE1 kinetics reflect both the large and the small frequencies albeit with different amplitudes; *second*, cAMP/PKA dynamics reflect only the small frequency of the calcium input; and *finally*, cAMP/PKA acts as a leaky integrator of calcium because of frequency attenuation by the intermediary steps (Figure 8). We also found that spines generated different spontaneous calcium frequencies in cultured neurons (Figure 6), thus motivating the need to study multiple time scales. These findings have implications for cAMP/PKA signaling in dendritic spines in particular and neuronal signal transduction in general.

**Figure 8:**
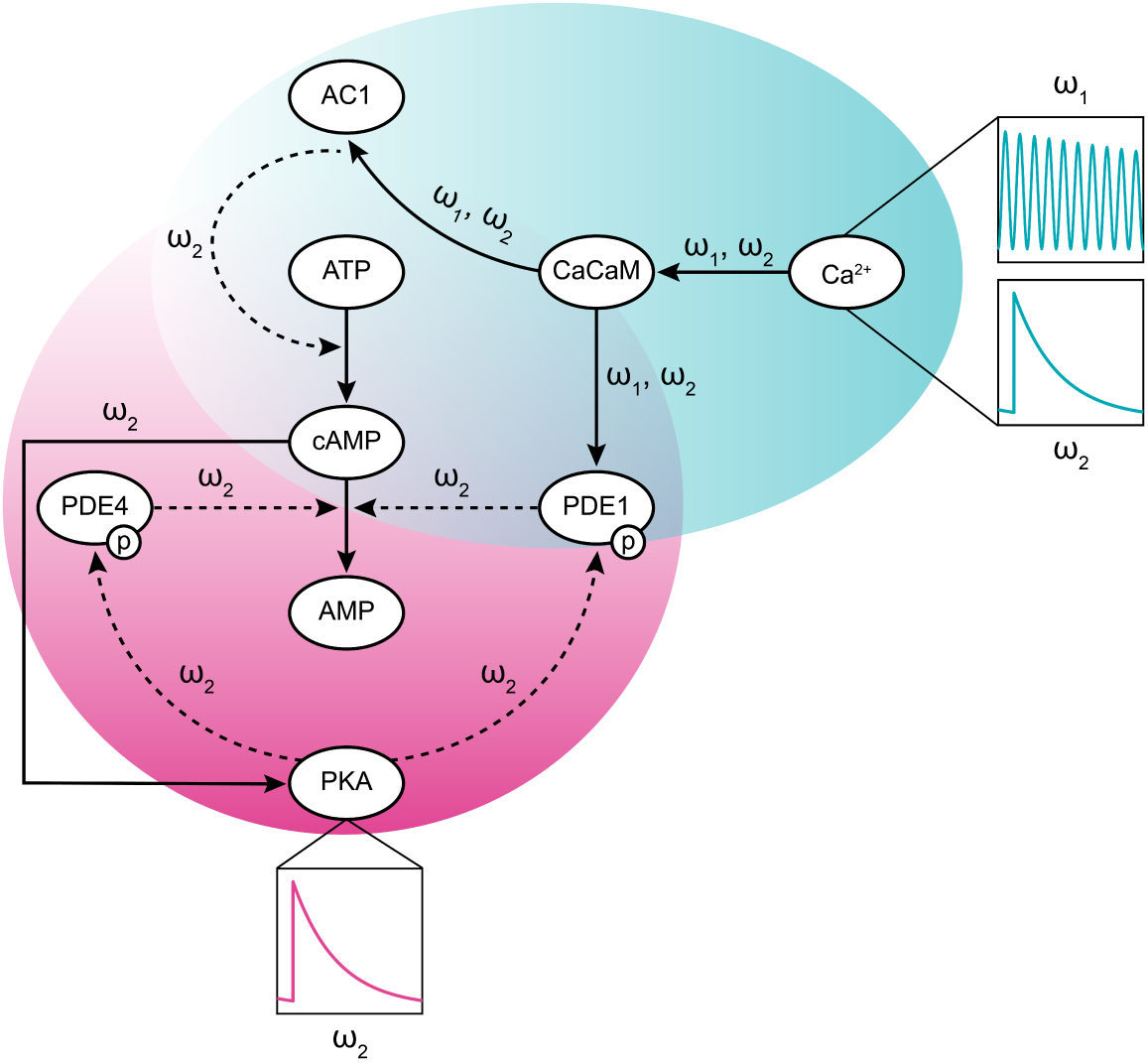
The frequency control in the modeled cAMP/PKA pathway from calcium stimulation through PKA. Calcium oscillates with a combination of frequencies (*ω*_1_=0.5 Hz and *ω*_2_=0.003 Hz). From calcium-calmodulin complex activation of AC1 and PDE1 enzymes through cAMP production and degradation, the effect of *ω*_1_ becomes trivial. Ultimately, through the downstream pathway, *ω*_2_ remains the only significant oscillatory frequency in this pathway.

The ability to modulate the output frequency in response to a signal input is an important feature in biochemical signal transduction; it ensures robustness and noise-filtering. As a result, the output signal, cAMP/PKA in this case, will respond to calcium input for strong signals but not to weak signals (Figure 7). In general, the combination of activation and inactivation kinetics along with feedback loops, many of which oscillate, may be responsible for noise attenuation in signal regulation [117, 118]. Noise attenuation has a significant role in regulating biological switches. These switches must sense changes in signal concentration while buffering against signal noise [119]. Our model predicts that depending on the sensitivity of regulating enzymes to Ca^2+^, the signaling input can be modified to the desired signaling output with a different range of frequency. For example, starting with a stimulus Ca^2+^ signal with a combination of *ω*_1_ and *ω*_2_ oscillation frequencies, the cAMP/PKA pathway can modify the initial signal and filter it so that components downstream in the pathway, such as PKA receive a signal with an oscillation frequency of *ω*_2_ (Figure 8).

As a result, cAMP/PKA can be thought of as a leaky integrator of calcium signals (Figure 5A and B). This signal modification is important in the context of how cAMP/PKA directly impacts spine morphology [120], receptor trafficking [121], and transcription factor activation (e.g. CREB and Epac) [32, 122]. While these transcription factors are not directly considered in our study, it is possible that the frequency modulation exhibited by the pathway we modeled plays a critical role in contributing to neuronal robustness to small fluctuations while retaining the ability to rapidly respond to large calcium influxes. In order to identify whether calcium or G_*s*_ signals dominate, we added the reactions pertaining to the Gs-mediated cAMP production [43] in our model. Interestingly, we found that the G_*s*_ pathway had very little effect on the conclusions of our model. The main reason for this is that calcium mediates both AC1 and PDE1 activation, thereby creating a strong incoherent feedforward loop for cAMP [123], whereas G_*s*_ linearly activates AC1 and PDE1 depends only on PKA feedback. Of course, the addition of other feedback loops and regulatory pathways in future efforts will dissect this crosstalk better.

The calcium input to our model was inspired by measurements of spontaneous calcium dynamics in dendritic spines (Figure 6). Ideally, we would like to use imaging to simultaneously measure spontaneous calcium and cAMP/PKA activity in dendritic spines in order to test our model predictions. However, despite the recent developments in fluorescence biosensing methods, imaging and co-imaging of multiple second messengers, protein kinases, and phosphatase activities in micron-sized compartments remains challenging [124]. This is also true for simultaneously imaging the dynamics of both cAMP and calcium in single dendritic spines [36, 125]. Recently, Tang *et al.* measured PKA activity in dendritic spines in response to glutamate uncaging using new FRET (fluorescence resonance energy transfer) probes that they developed [36]. The timescale of PKA dynamics from our model (Figure 7A) are in good agreement with the time scale of approximately 10 min (Figure 6 of Tang *et al.* [36]). This indicates our developed model closely mimics reported PKA activity in subcompartments, in this case, dendritic spines.

Despite the predictive capability of our model, there are certain limitations that must be acknowledged. First, we only consider a deterministic model and have currently ignored stochastic dynamics in the system. It is possible that the frequency modulation effects presented here will be somewhat altered by including stochasticity but we anticipate that the overall conclusions will not be affected on average if the same network is considered. An additional limitation of our work emerges in the form of challenges associated with co-imaging two second messengers simultaneously in the same spine. We are working on developing such probes to overcome these challenges. Finally, there are indeed many protein kinases and phosphatases that affect this pathway [126–129] and ongoing efforts in our group are focused on building up increasingly comprehensive network maps of these signaling reactions. Furthermore, spatial regulations of cAMP signaling are other important factors that need to be considered [130–132]. Enzyme regulation seems to play a significant role in cAMP/PKA pathway dynamics and signal filtration [133–135]. These enzymes can also be localized by A-kinase anchoring proteins (AKAPs) and change cAMP/PKA dynamics further [136–138]. The effect of spine size and enzyme localizations are considered in a companion study [62].

## Author Contributions

DO and PR conceived the study; DO developed the model and conducted the simulations; DLS conducted neuronal culture experiments; BC aided in experimental design; all authors analyzed the data and wrote the manuscript.

## Acknowledgments

The authors wish to thank Miriam Bell, Justin Laughlin, and Allen Leung for their valuable comments. This work was supported by AFOSR MURI grant number FA9550-18-1-0051. DLS was supported by NIH/NCI T32 CA009523. Confocal imaging was done at the UC San Diego Nikon Imaging Center with the assistance of Eric Griffis. DLS would like to thank AJ Slepain for her assistance with neuronal culture.

