## Supplemental Information for "Computational modeling reveals frequency modulation of calcium-cAMP/PKA pathway in dendritic spines"

### Reaction kinetic type selection for PDE1 activation

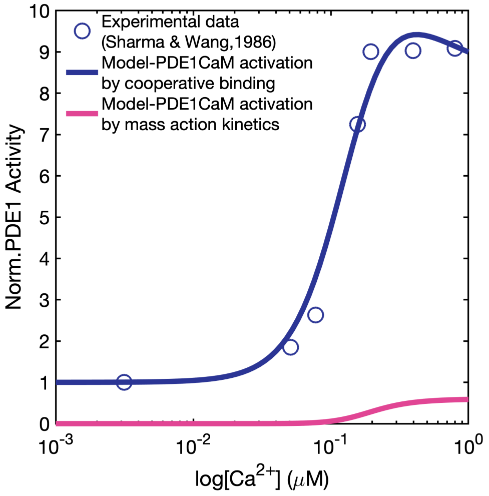


Figure S1: Cooperative binding kinetics shows a better fit for PDE1 activation by $Ca^{2+}$ in comparison to mass-action kinetics.

### Stochastic vs Deterministic Simulations

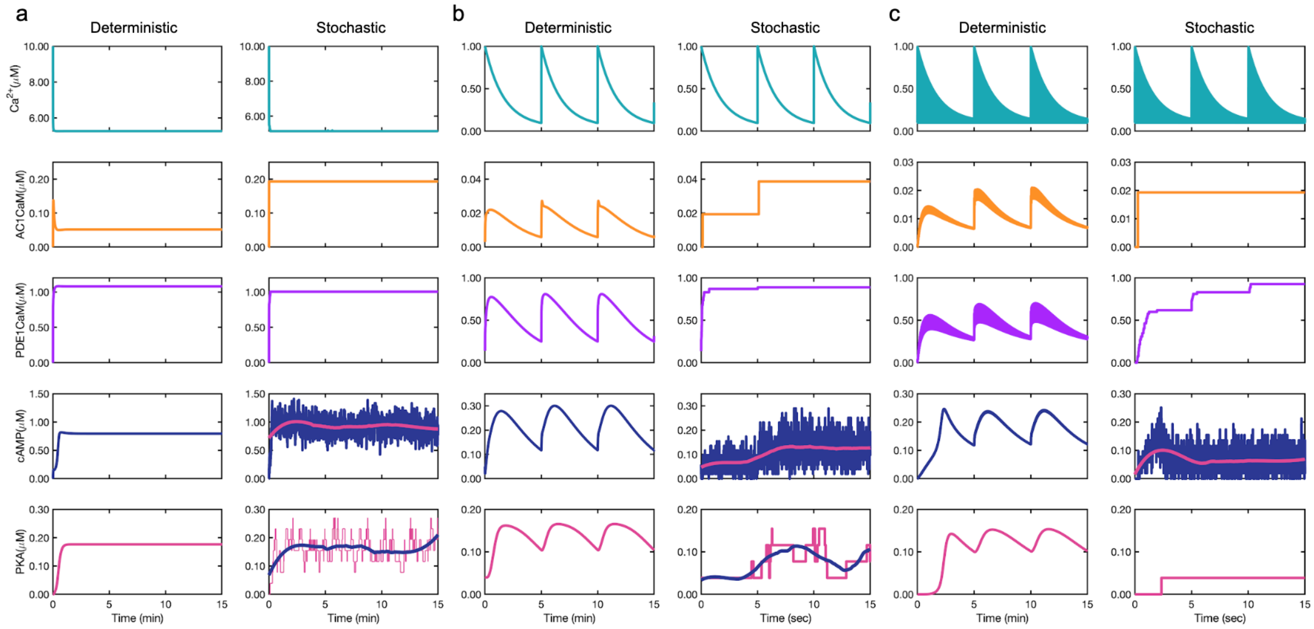


Figure S2: Comparison of stochastic and deterministic simulations for a) Non-oscillating $Ca^{2+}$, b) $Ca^{2+}$ oscillating in minute scale, and c) $Ca^{2+}$ oscillating in second scale.

### Effect of AC1 activation by $G_s$ on cAMP production

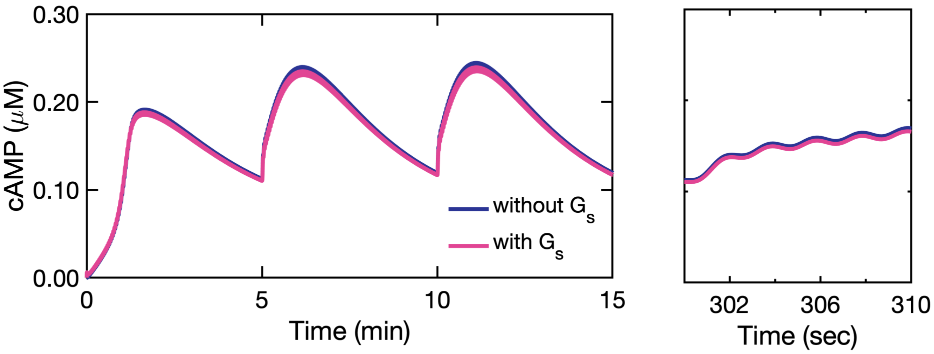


Figure S3: AC1 activation by $G_s$ does not affect cAMP production.

### Other oscillation frequencies

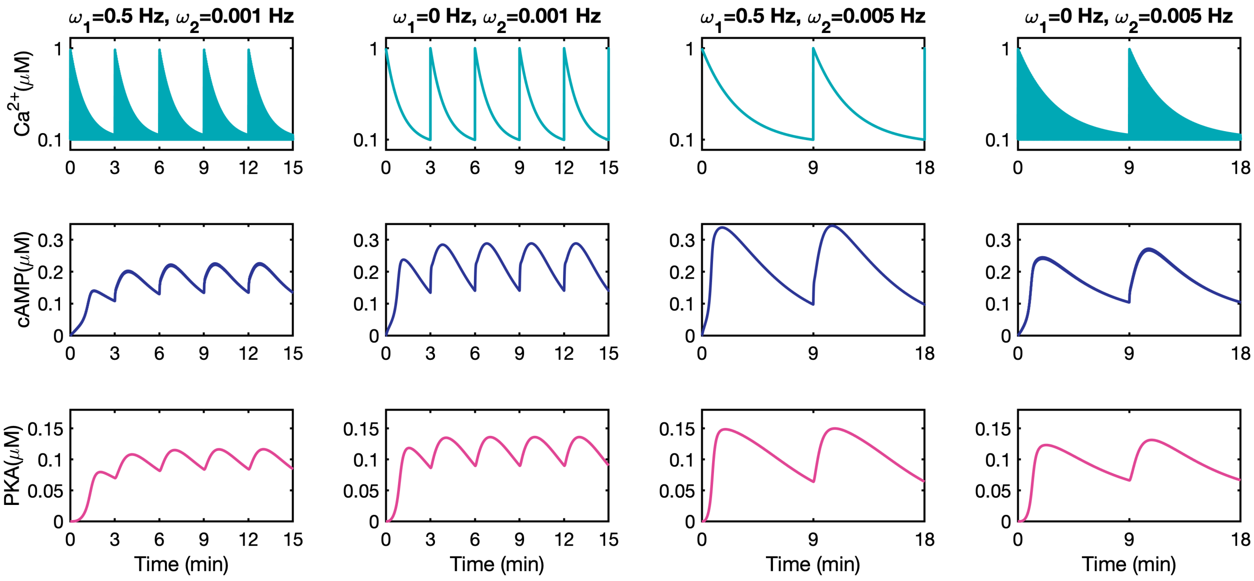


Figure S4: Two other calcium minute-scale frequencies ($\omega_2$=0.001 and $\omega_2$=0.005) along with second-scale oscillations ($\omega_1$=0.5) were chosen to confirm that cAMP and PKA only pick up minute-scale oscillations of calcium.
